# Biofilm formation displays intrinsic offensive and defensive features of *Bacillus cereus*

**DOI:** 10.1101/676403

**Authors:** Caro-Astorga Joaquin, Frenzel Elrike, Perkins James Richard, Antonio de Vicente, Juan A.G. Ranea, Oscar P. Kuipers, Romero Diego

## Abstract

**Background:** Biofilm formation is a strategy of many bacterial species to adapt to a variety of stresses and has become a part of infections, contaminations or beneficial interactions. We previously observed that *B. cereus* ATCC 14579 (CECT148), formed a thick biomass of cells firmly adhered to abiotic surfaces.

**Results:** In this study, we combined two techniques, RNAseq and iTRAQ mass spectrometry, to demonstrate the profound physiological changes that permit *Bacillus cereus* to switch from a floating to a sessile lifestyle, to undergo further maturation of the biofilm, and to differentiate into offensive or defensive populations. The rearrangement of nucleotides, sugars, amino acids and energy metabolism lead to changes promoting reinforcement of the cell wall, activation of ROS detoxification strategies or secondary metabolite production, all oriented to defend biofilm cells from external aggressions. However, floating cells maintain a fermentative metabolic status along with a higher aggressiveness against hosts, evidenced by the production of toxins and other virulent factors.

**Conclusions:** We show that biofilm-associated cells seem to direct the energy to the individual and global defense against external aggressions and competitors. By contrary, floating cells are more aggressive against hosts. The maintenance of the two distinct subpopulations is an effective strategy to face changeable environmental conditions found in the life cycle of *B. cereus*.

## Background

*Bacillus cereus* is a widespread bacterium that can colonize a multitude of niches, including soil and seawater, where it survives living as a saprophyte or in transit from other ecological niches. This bacterium can also be found in association with plant tissues, living as a commensal or in symbiosis as a rhizosphere inhabitant [1]. Mammalian and arthropod guts are also a niche for *B. cereus*, where it can live as a commensal or a pathogen, either opportunistic or not [2, 3]. The versatility shown by these bacteria evidence the resilience ability against a wide range of environmental conditions [4]. *B. cereus* gives its name to the *B. cereus sensu lato* group, which includes the phylogenetically similar bacterial species *B. thuringiensis* and *B. anthracis*, diverse in their host impact as they affect insects and human health [5]. Some strains of *B*. cereus provide benefits to plants, as a promoter of growth (PGP) or biocontrol against microbial diseases, while others are proposed as probiotics for cattle and even for humans [6, 7]. By contrast, there are strains responsible for human pathologies mainly caused by food poisoning, contamination of products in the food industry or even food spoilage [8–10]. Biofouling, clogging and corrosive consequences of *B. cereus* in industrial devices complete the concerns of humans regarding this bacteria species [10].

Regardless of the consequences, most of the scenarios listed above are believed to be related with the organization of bacterial cells in biofilms. The formation of biofilms is considered an important step in the life cycle of most bacterial species, and it is known to be related to outbreaks of diseases, resistance to antimicrobials, or contamination of medical and industrial devices [11]. Approximately 65% of bacterial human diseases are estimated to involve bacterial biofilms, thus these multicellular structures might be considered potential targets to fight against bacterial diseases [12]. Based on the relevance of bacterial biofilms, our research focuses on elucidating the intrinsic factors employed by *B. cereus* to switch to this sedentary lifestyle. In general, it is known that after encountering an adequate surface, motile bacterial cells switch from a floating or planktonic to a sessile lifestyle followed by the assembly of an extracellular matrix. Studies on biofilm formation in the Gram-positive bacterium *Bacillus subtilis* have substantially contributed to our understanding of the intricate machinery devoted to efficiently complete this transition [13]. While studies on biofilm formation on specific *B. cereus* strains indicate that key processes resemble *B. subtilis* biofilm development, clear differences start to be perceived, representative of the evolutionary distance between the two species [14]: i) the minor role of the exopolysaccharide of *B. cereus* homologous to the *epsA-O* of *B. subtilis* in biofilm formation [15]; ii) the absence of homologues to the accessory protein TapA, necessary for amyloid-like fiber assembly in *B. subtilis*, iii) the existence of two paralogs of *subtilis* TasA, i.e., TasA and CalY [16]; iv) the absence of the hydrophobic BlsA protein, which coats the biofilm in *B. subtilis* and play a role in the biofilm architecture [17]; v) the differences in the regulatory networks of biofilm formation, lacking the regulatory subnetworks II and III that involve SlrA-SlrR-SinR and Abh; and the gain of the pleiotropic regulator PlcR involved in virulence and biofilm formation [14, 18]; vii) the absence in *B. cereus* of the lipoprotein Med associated with KinD phosphorylation activity that triggers biofilm formation; and viii) the different adhesive properties of the spores of *B. cereus* [19]. Therefore, novel in-depth studies are required not only to confirm similarities between the two microbes but also to highlight unexplored differences.

In a previous work, we reported that the growth of *B. cereus* ATCC 14579 (CECT148) biomass of cells adhered to abiotic surfaces is a process that clearly increases with time [16]. A genomic region containing the two paralogous proteins TasA and CalY, the signal peptidase SipW and the locus *BC_1280* were proven essential in the transition from planktonic or floating to sedentary and further growth of the biofilm. The differences found in *B. subtilis* in this and other reports led us to investigate which are the additional intrinsic genetic features that warrant *B. cereus* to solve hypothetical environmental situations by the assembly of biofilms. The combination of two techniques, RNA sequencing (RNAseq) and mass spectrometry proteomic (isobaric tags for relative and absolute quantitation – iTRAQ), enabled us to acquire solid evidence of the global changes differentiating floating from biofilm programmed cells and depict how biofilm of *B. cereus* progresses. We report the reinforcement of the cell wall of biofilm cells, that would prepare cells for further assembly of macromolecules as polysaccharides and other adhesins, and additional protection of cells individually from external aggressions; and the major production of secondary metabolites of biofilm-associated cells to defend against competitors. Additionally, floating cells are maintained in a sustained stationary phase of growth conducive to survival, not in the form of spores, and more aggressive against the human host. Our findings argue in favor of the metabolic versatility of *B. cereus* and the fine tuning in gene expression which permit this specie to maintain the two distinct subpopulations necessary to face changeable environmental conditions.

## Results and discussion

### Massive changes in gene expression define different stages in biofilm formation

To study the transition of cells from floating to biofilm and its further progression, we used static cultures and developed the experimental setup summarized in Figure S1 (described in detail in the section Methods). Regarding biofilm formation as a developmental program, we considered 24 h cultures as the initial time to perform the comparison, when biomass started to be visible and thus suitable for collection and further analysis. This comparison allowed us to define the initial changes in gene expression which characterize sessile or floating cells in young biofilms. This comparison was used as a reference and a further time-course analysis was made to study the changes arising at 48 h, when the biofilm is consolidated in thickness and adherence, and at 72 h, when the biofilm is fully mature (Fig. S2A) [16].

Although transcription and translation are intimately associated in bacteria in space and time, there are other factors that can alter the final amount of protein and, subsequently, the final function trusted in those proteins. In *B. cereus*, it has been reported that mRNA shows a half-life ranging from 1-15 minutes [20]. Several studies on *Bacillus* species classify the total amount of proteins inside a cell into two protein populations of either a labile fraction or a fraction of long-lived proteins with a turnover ranging from less than one hour to forty hours, which can introduce large differences between mRNA transcripts and protein function [21]. Therefore, our samples were analyzed by two complementary quantitative techniques directed to two steps in gene expression: i) sequencing total mRNA analysis and ii) quantitative proteome analysis using iTRAQ. Like RNAseq, iTRAQ is very accurate in defining quantitative differences, although it requires a relatively large amount of protein. Thus, RNAseq data were considered as a reference in our study for gene expression, and the iTRAQ data were used to confirm that variations in gene expression result in the same direction as variations of translation. Henceforth, confirmed bacterial factors will be indicated with an asterisk “*” and discordances with a “x”. The absence of a mark means that this protein was not detected by iTRAQ.

The genome of *B. cereus* ATCC 14579 possesses 5490 open reading frames (ORFs), and the RNAseq analysis showed that 1292 genes were differentially expressed (log2<>|2|) in biofilm-associated cells compared to floating cells, meaning 23,5% of the total gene content. These numbers are illustrative of the outstanding genetic machinery dedicated to the developmental program that leads to the formation of a bacterial biofilm (Fig. S2B, C). Proteomic analysis revealed 945 proteins with differential expression (log2>|0.7|) at any of the times sampled. A wider view of the data indicates a good correlation between both techniques in the number of genes with up or down expression levels (Fig. 1A). Although the initial putative suspicion on the relative low concordance of transcription and translation, discordances among mRNA and protein levels were limited to only 43, 9 and 21 hits at 24, 48 and 72 h respectively, a finding that shows a first confirmation of data.

**Fig 1.**
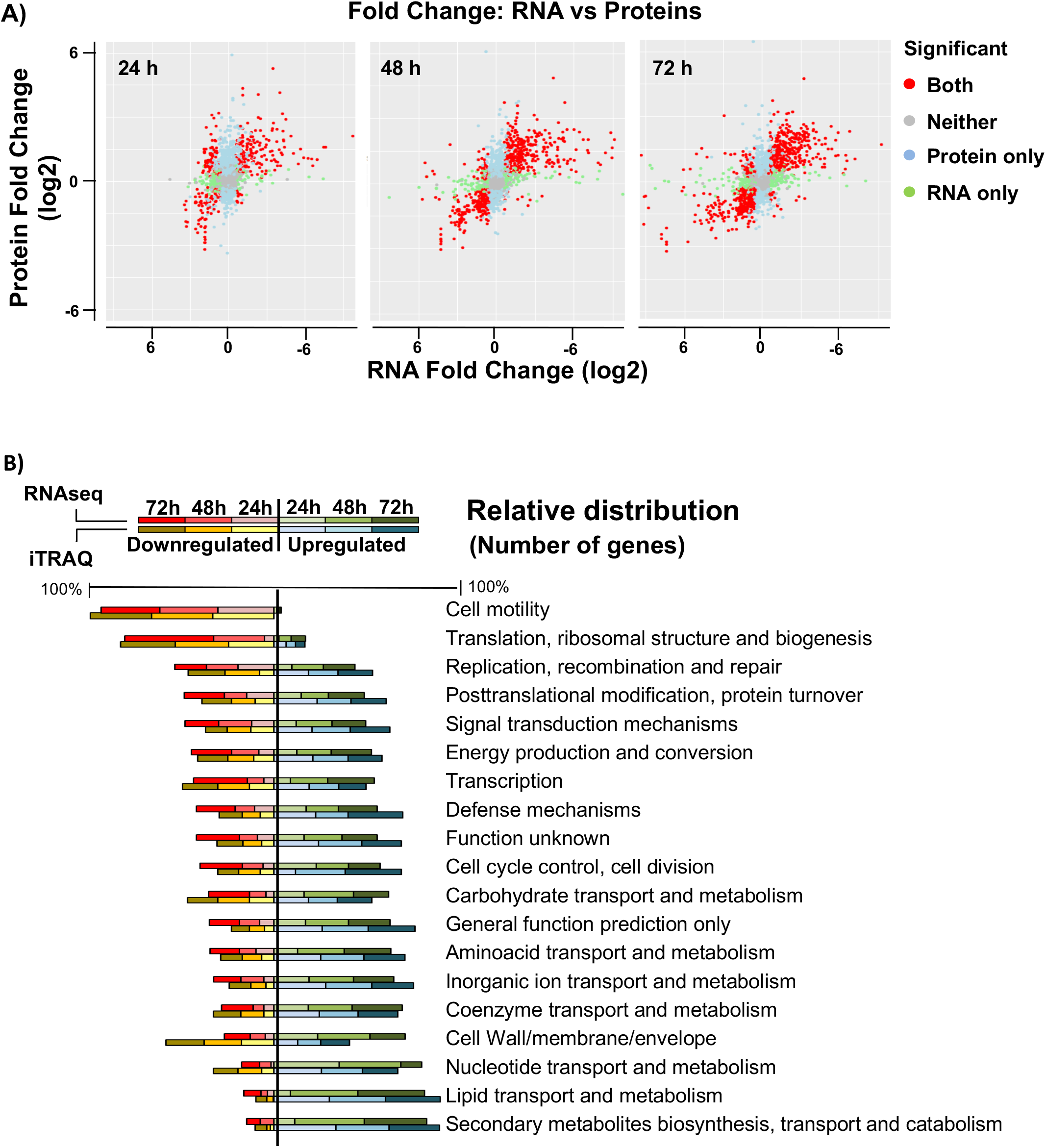
Validation of variation of gene expression using RNAseq and iTRAQ analyses. **A)** Fold change in expression between transcriptomic (RNA-Seq) and proteomic (iTRAQ) results from samples at 24, 48 and 72 h. Points show significant changes (adjusted p-value < 0.05) according to both technologies (red), changes only found in proteomic (blue) or only found in transcriptomic (green). Grey dots represent items detected with statistically no significant changes. The top right and bottom left quadrants show concordant directions of change between both technologies; the top left and bottom right show conflicting directions. **B)** Relative comparison of COG categories of genes and proteins. Total number of elements of each category were relativized to a percentage and distributed in down or upregulated. This graph shows that the ratio of elements that are up or downregulated within each COG category is conserved, revealing similar results in terms of functional categories among RNAseq and iTRAQ techniques.

To obtain a detailed view of the major physiological changes, genes with differential expression were sorted into Cluster of Orthologous Groups (COG) categories (Fig. 1B). The data showed some foreseen changes such as the downregulation of the COG for flagellum assembly compared to that of floating cells. Delving into more detail, as COG categories ignore many of the individual differentially expressed ORFs, manual classification of the 1292 genes into functional groups was performed. This manual classification also showed an expected behavior in *B. cereus* in terms of sporulation, which group of genes are upregulated as it is linked to biofilm formation. In addition, some known regulators of sporulation, such as the positive biofilm regulator SinI [22], are upregulated, while the negative biofilm regulator PlcR, which is negatively controlled by Spo0A-P [23], is downregulated.

We made an analysis to look for time-specific function changes but most of the functions show a more pronounced up or downregulation over time. At 24 h, genes of pyrimidine metabolism are specifically upregulated in biofilm cells (Table 1). The clearest time-specific change affect sporulation. Clustering of the 124 sporulation-related genes by expression pattern using the Short Time-series Expression Miner (STEM) showed an increase of expression at 24 h and 48 h and repression at 72 h of biofilm formation (Fig. S4A). Cluster 6 of the STEM clustering results (Fig. S4B) showed a small group of genes with an overexpression pattern at 72 h compared to previous stages, but these genes are specific negative sporulation regulators. Despite the arrest in the expression of sporulation genes, the sporulation process inertia reveals a 9% of spores within the 72h-biofilm (Fig. S9).

**Table 1.** Genes of the pyrimidine metabolism specifically upregulated in biofilm cells. Asterisk indicates confirmed behaviour at protein levels.

| Log2 (fold change) |  |  |  |  |  |
| --- | --- | --- | --- | --- | --- |
| Gene ID | 24h | 48h | 72h | Function. Gene name | iTRAQ |
| **BC0325** | 2.41 | 2.56 | -4.27 | Purine biosynthesis pathway. Adenylosuccinate lyase | * |
| **BC0326** | 3.13 | 3.87 | -0.69 | Purine biosynthesis pathway. Phosphoribosylaminoimidazole-succinocarboxamide synthase | * |
| **BC0327** | 3.09 | 3.37 | -0.66 | Purine biosynthesis pathway. Phosphoribosylformylglycinamide synthase subunit PurS | * |
| **BC0328** | 2.91 | 3.37 | -0.80 | Purine biosynthesis pathway. Phosphoribosylformylglycinamide synthase I | * |
| **BC0329** | 2.37 | 3.17 | -1.16 | Purine biosynthesis pathway. Phosphoribosylformylglycinamide synthase II | * |
| **BC0330** | 2.64 | 3.91 | 0.40 | Purine biosynthesis pathway. Amidophosphoribosyltransferase | * |
| **BC0331** | 2.96 | 3.52 | -1.96 | Purine biosynthesis pathway. Phosphoribosylaminoimidazole synthetase | * |
| **BC0332** | 4.01 | 3.68 | -1.38 | Purine biosynthesis pathway. Phosphoribosylglycinamide formyltransferase | * |
| **BC3882** | 3.66 | -0.38 | -0.07 | Pyrimidine biosynthesis pathway. Orotate phosphoribosyltransferase | * |
| **BC3883** | 3.73 | 0.65 | 0.84 | Pyrimidine biosynthesis pathway. Orotidine 5'-phosphate decarboxylase |  |
| **BC3884** | 4.14 | 1.06 | 1.54 | Pyrimidine biosynthesis pathway. Dihydroorotate dehydrogenase 1B |  |
| **BC3885** | 4.18 | 0.98 | 1.12 | Pyrimidine biosynthesis pathway. Dihydroorotate dehydrogenase electron transfer subunit |  |
| **BC3886** | 4.29 | 0.22 | 0.95 | Pyrimidine biosynthesis pathway. Carbamoyl phosphate synthase large subunit | * |
| **BC3887** | 3.40 | 1.19 | 0.66 | Pyrimidine biosynthesis pathway. Carbamoyl phosphate synthase small subunit |  |
| **BC3888** | 5.69 | 2.74 | 2.27 | Pyrimidine biosynthesis pathway. Dihydroorotase |  |
| **BC3889** | 4.87 | 1.78 | 0.26 | Pyrimidine biosynthesis pathway. Aspartate carbamoyltransferase |  |

These time-resolved analyses indicate that most of the biofilm functions are intrinsically associated with the biofilm state, with only some exceptions of processes only required in specific developmental stages.

### Changes in the metabolic activity of biofilm cells to satisfy the synthesis of the new extracellular matrix

It appears that cells committed to the biofilm lifestyle have specialized in certain metabolic pathways most likely directed to the synthesis of proteins, polysaccharides and extracellular DNA (eDNA), elements of the extracellular matrix necessary to cover a range of functions from adhesion to biofilm architecture [17, 24, 25]. The extracellular matrix plays an important role in defense against physical stress like desiccation [26]. In our results, we found overexpression of multiple genes involved in the synthesis of the extracellular matrix (Fig. 2A). Our analysis revealed that the well-known structural biofilm-related genes *sipW, tasA* and *calY* (*BC1278*, BC1279*, BC1281\**) are overexpressed in biofilm-associated cells. Not surprisingly, and in agreement with findings from *B. subtilis* biofilm studies, the master regulator that controls the expression of these genes, SinR, and its repressor SinI [27], were upregulated in biofilm-associated cells. In a previous work, the genes *purA* (*BC5468*), *purC* (*BC0326*) and *purL* (*BC0327*), which are implicated in purine biosynthesis, were demonstrated to be indispensable for biofilm formation in *B. cereus* [28–30]. In our analysis biofilm cells upregulate the entire gene cassettes for pyrimidine (*BC3883-89*) and purine (*BC0325-BC0332\**) biosynthesis, as well as transporters of amino acid and formate synthesis required for nucleoside synthesis (BC0403*, BC0639, BC0640*, *BC2383*, BC3939*, BC2790), nucleoside transporters (*BC2973, BC0363*) and specific phosphate transporters (*BC4265-68* and *BC0711*), all molecules required for DNA synthesis. These results suggest that these metabolic changes might be, among others, oriented to the synthesis of the eDNA of the extracellular matrix.

**Fig. 2.**
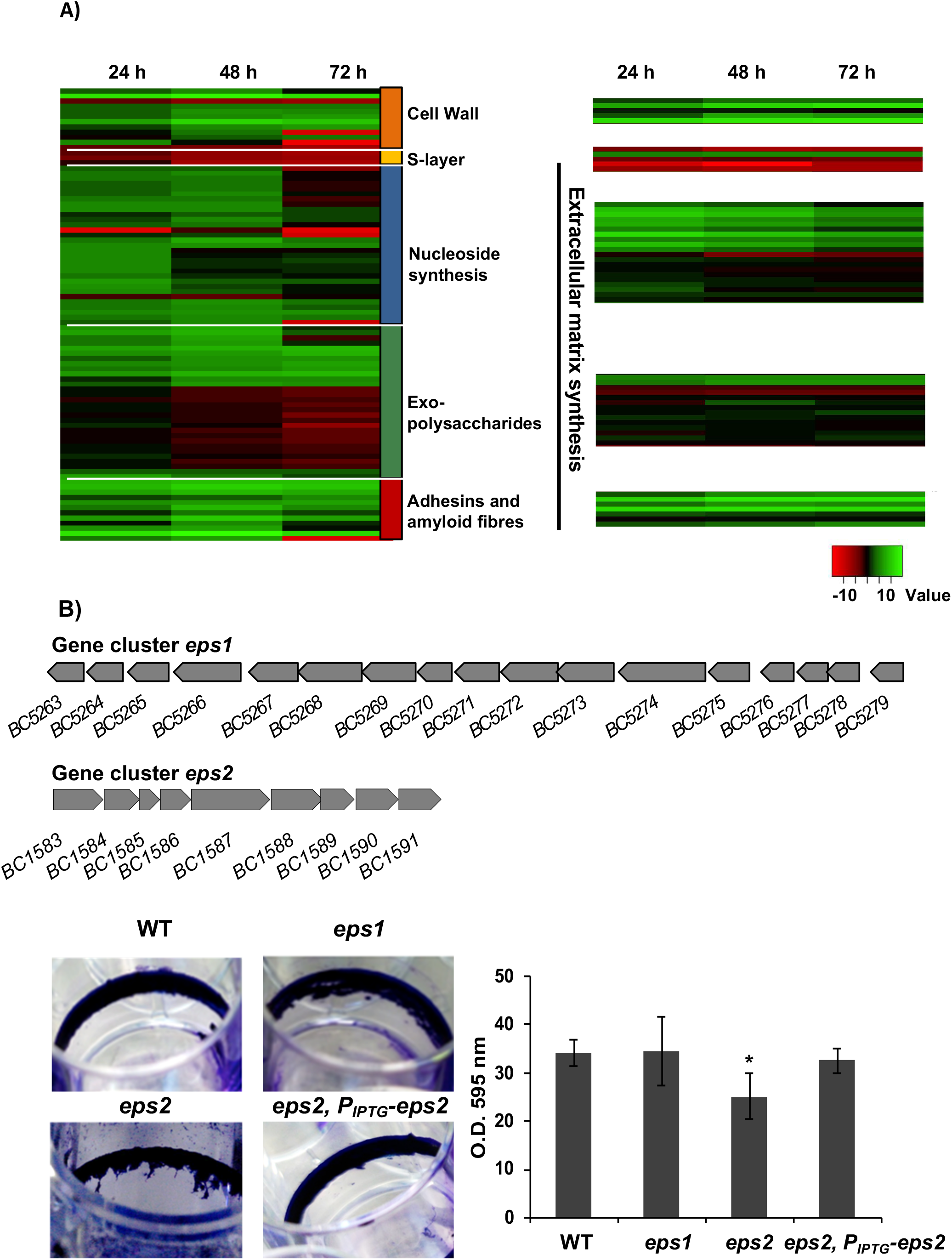
Expression dynamics of external elements and extracellular matrix components during biofilm formation. **A)** Expression dynamics of external elements and extracellular matrix components during biofilm formation. Comparison of heat maps from RNA-seq (left) and iTRAQ (right) results at 24, 48 and 72 h, over a selection of elements of the outer layers of *B. cereus* cells and components of the extracellular-matrix shows the following: i) The low expression levels of S-layer components, the major relevance of eDNA at 24 h-48 h, and the continuous overexpression of factors related to cell wall synthesis, some exopolysaccharides and other adhesins. **B)** Gene clusters for the synthesis of two hypothetical exopolysaccharides: top, *eps1*, homologous to the *eps* region in *B. subtilis* and bottom, *eps2* specific of *B. cereus*. Crystal violet staining of adhered biofilms showed no differences between *eps1* single mutant and wild type, and slight reduction of biofilm formation of *eps2* single mutant. Reversion of crystal violet staining in the strain *eps2* complemented with a replicative plasmid (pUTE973) harboring the transcriptional construction P_IPTG_-eps2. Pictures were taken 72 h after growing cells with no agitation at 28°C in TY. The adhesion to abiotic surface of the different strains in TyJ medium at 48 h was measured by the amount of crystal violet retained in the bacterial biomass. (error bars correspond to SD.*, differences statistically significant, p□0.05, T-student, 3 replicates).

In addition to proteins and eDNA, exopolysaccharides (EPSs) constitute an integral component of the extracellular matrix of most bacterial biofilms. Importantly, our data indicate noticeable differences in the well-defined role of EPS in *B. subtilis*. First, the putative EPS gene cluster *BC5263-BC5279\** (Fig. 2B, top) homologous to the *eps* operon of *B. subtilis*, showed a similar expression pattern to that of floating cells. Indeed, knockout mutants in this genetic region (herein *eps1*) were not arrested in biofilm formation based on the crystal violet staining of the adhered biomass (Fig. 2B, bottom). This finding is in agreement with previous studies showing that mutants in this gene cluster in *B. cereus* 905 did not affect biofilm formation [19]. Second, we identified an additional gene cluster (herein *eps2*), annotated as a capsular polysaccharide synthesis, which is overexpressed in biofilm-associated cells (*BC1583-BC1591*) (Fig. 2B, top). In agreement with a previous report, we could not microscopically visualize a capsule in this strain of *B. cereus*, which led us to propose a role of this new gene cluster in biofilm formation. A mutant in this region was not completely arrested but was partially impaired in the biofilm adhered to the wells (Fig. 2B, bottom). This deviation of phenotype was reversed by expressing the entire *eps2* region in trans, in a replicative plasmid under control of an IPTG inducible promoter (*eps2, PIPTG-eps2*). Attending to these results, we hypothesize that *eps2* region play a role in biofilm formation.

The biofilm architecture is very resource-consuming. The supply of amino acids might come from protein recycling or being compensated by downregulation of the synthesis of other polymers such as flagellum assembly, but this supply seems to be insufficient, according to our results. We found a noticeable upregulation of genes encoding transporters of specific amino acids (Supplemental Table S1), especially those genes encoding a glutamine transporter and a glutamine-binding protein, which were significantly upregulated in comparison to other specific amino acid transporters. Glutamine is an important amino acid in bacterial physiology, and beyond the well-known involvement in the synthesis of proteins, this amino acid serves additional functions as a growth control molecule through dynamic gradients of glutamine concentration along the colony. This gradient induces and reduces the metabolic activity of cells to allow nutrients to diffuse to the inner part of the biofilm. [32, 33]. Apart from this, the glutamine supply is consumed for important functions, such as the synthesis of the protein component of the extracellular matrix or cell wall poly-linking. The synthesis of amino acids is under the control of DnaK (*BC4858*), a suppressor protein homologous to DksA in *E. coli*, which is a negative regulator of rRNA general expression during starvation and a positive regulator of several amino acid biosynthesis, amplifying the effects of the secondary messenger ppGpp. In *E.coli*, DksA is present at similar concentrations at mid-log phase and in late stationary phase with unchangeable levels; however, the levels during biofilm formation were not explored [35]. In a biofilm of *B. cereus*, levels of DnaK are remarkably upregulated compared with that of floating cells. This led us to speculate that DnaK belongs to the complex biofilm regulatory network that participates in the coordination and synchronization of the protein synthesis machinery required for biofilm formation in *B. cereus*.

### Biofilm-associated cells thicken their cell wall

In addition to the collective protection to antimicrobials that can be provided by the extracellular matrix, our data suggest that individual cells might develop protection against external aggressors by enhancing the physical barrier represented by the cell wall. Besides, in *B. subtilis*, it has been reported the importance of transglycosylation and transpeptidation of the PG, as well as the wall teichoic acid for biofilm formation, which disturbance affect the cell wall structure and drastically affect the extracellular matrix anchoring [36, 37]. Lipoteichoic acid (LTA) is a major component of the cell wall in Gram-positive bacteria, and the LTA synthase (*BC3765*) is upregulated in 48- and 72-hour aged biofilms. Peptidoglycan (PG), the other main component of the cell wall, is composed of units of N-acetylglucosaminic acid and N-acetylmuramic acid. In our experiments, we found that cell wall degradation enzymes are downregulated (*BC0888*, BC5237, BC3257, BC1660, BC3677*), except for the N-acetylmuramoyl-L-alanine amidase (*BC2823\**) which is specifically upregulated in biofilm cells. In addition, several penicillin-binding proteins (*BC1469, BC2190*, BC2448, BC2688, BC4075*), which are involved in the final stages of PG synthesis, are upregulated in biofilms with a maximum expression level at 48 h, as well as three transpeptidases (*BC2468*, BC5027, BC0771*) which catalyze the PG cross-linking (Fig. 3A). D-alanine is an important component of the cell wall which is implicated not only in the peptide cross-linking but also in the D-alanylation of teichoic acid, a modification reported to contribute to the resistance to several antimicrobials in *B. cereus* [38], and to biofilm formation, host defense or mouse intestinal tract colonization in other Gram-positive bacteria [39–42]. The bidirectional conversion of L-alanine into D-alanine is catalyzed by alanine racemases, and *BC0264* is upregulated at 24 h and 48 h. These gene are orthologous to *Dal-1* in *B. anthracis*, which is known to rescue the phenotype of a D-ala auxotrophic strain of *E. coli* [43]. These findings on the upregulation of the enzymatic machinery involved in PG, LTA and D-alanine synthesis in conjunction with the downregulation of enzymes implicated in cell wall degradation in biofilm cells, raised the question of whether there is an alteration in the cell wall thickness of biofilm-associated cells compared to floating cells. To explore this hypothesis, we measured the cell wall thickness of cell sections of 48-h biofilm and floating cells by electron microscopy, revealing a 33.4% increase in the cell wall thickness of biofilm cells, a structural change which was also observed in sporulating cells (Fig. 3C).

**Fig 3.**
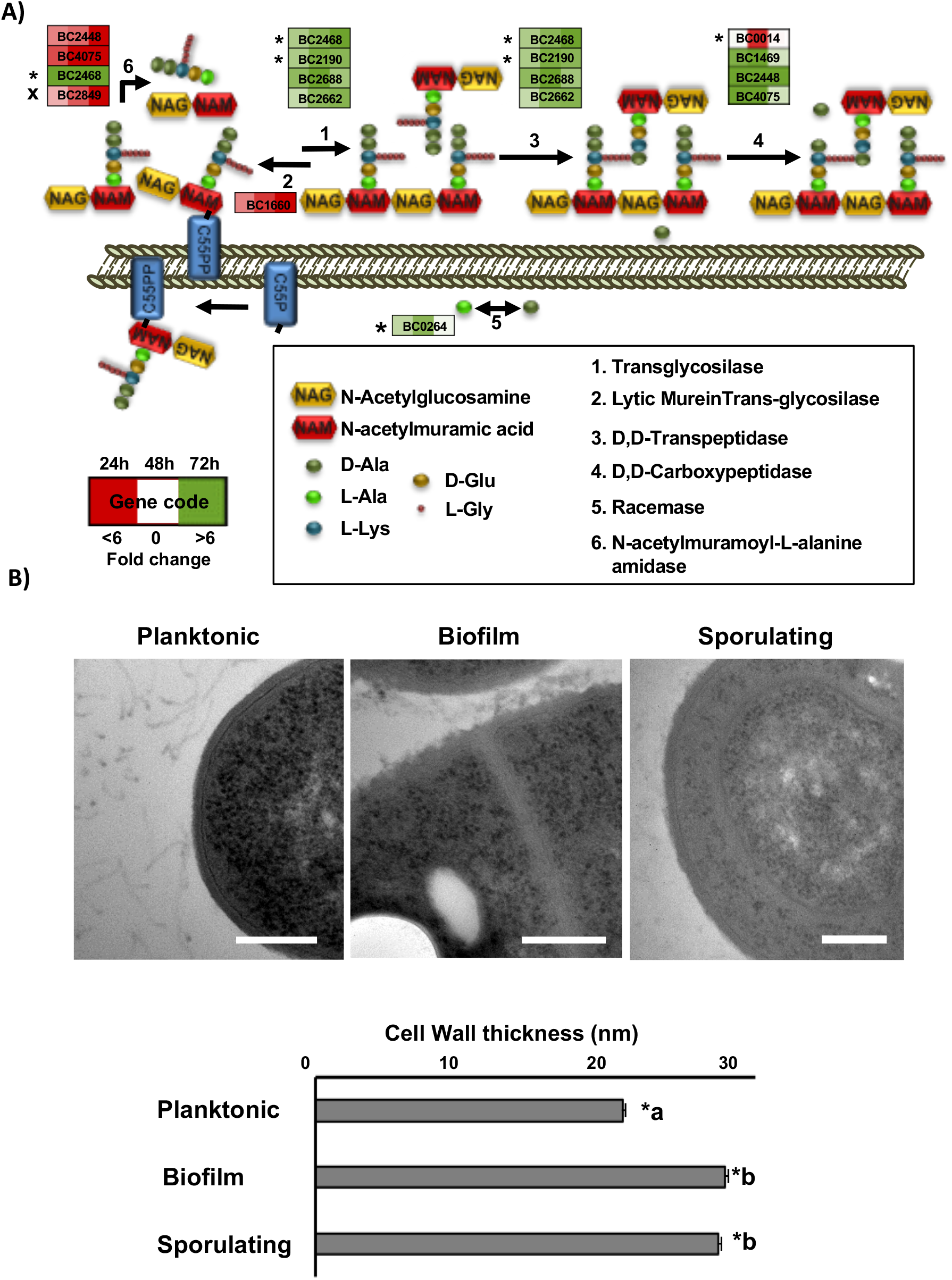
Cells in a biofilm increase the thickness of the cell wall. **A)** Scheme of the extra cytoplasmic cell wall synthesis pathway. Each step is performed by several genes represented in boxes. Colored squares inside boxes represent 24, 48 and 72 h sampling times, and the intensity of color shows the level of upregulation (green) or downregulation (red). Stars indicate confirmation of the behavior by iTRAQ results and cross indicate conflicting results. **B)** Transmission electron micrographs of floating cells, biofilm cells and sporulating cells inside the biofilm structure. Measurements of the cell wall thickness was done with ImageJ Fiji software over 40 images and 300 measures around the edges of each cell, showing an increase of 33.4% in the thickness of biofilm and sporulating cells. Standard deviation SC<0.01. Bars equal 200 nm.

### Biofilm-associated cells produce higher amount of secondary metabolites

Physical stresses are mainly buffered by the extracellular matrix, but how can bacteria defend from attacks from other bacteria in the competition for the space in the soil, plant rhizosphere or human gut? One way in which sessile bacterial cells may efficiently fight competitors regards the offense inflicted by antimicrobials. *B. cereus* possesses several genomic regions dedicated to the production of secondary metabolites with antimicrobial activity. In addition to these, we found genes with unknown functions that after *in silico* analysis (see the section Methods) appeared to be hypothetically involved in the production of putative secondary metabolites. Interestingly, all of them are upregulated in biofilm-associated cells (Supplemental Table S2). Among those regions were genes encoding NRPS/PKS or putative bacteriocin-synthesizing proteins such as i) thiocillin, ii) tylosin-like, iii) bacitracin, iv) porphyrins, v) a putative member of the heterocycloanthracin/sonorensin family of bacteriocins; vi) other putative bacteriocin (*BC1248-50*) vii) the cluster *BC1210-BC1212* orthologous to a streptomycin synthesis gene cluster in *B. subtilis;* and viii) a colicin-like toxin. Colicin is a bacteriocin which has been proposed to play an important role in microbial competition in the gut and to facilitate intestinal colonization [44]. There is a gap in the study of secondary metabolites in *B. cereus* to confirm that these genomic regions are responsible for the synthesis of active compounds. Only thiocillin has been studied in detail and has been proven to produce an active peptide. As the iTRAQ method is unable to detect these peptides and given that this antimicrobial is presumably produced in higher amounts by biofilm cells, we analyzed the presence of this metabolite in floating and biofilm cells fractions using mass spectrometry. This analysis confirmed that thiocillin was present in the pellet and supernatant of biofilm cells, but below detectable levels in the pellets of floating cells and in the spent medium (Fig. 4 and supplemental Fig. S5). Based on this observation, one might speculate that the production of these antimicrobials is used to colonize a niche and for protection against competitors which are in close contact in the direct surroundings.

**Fig 4.**
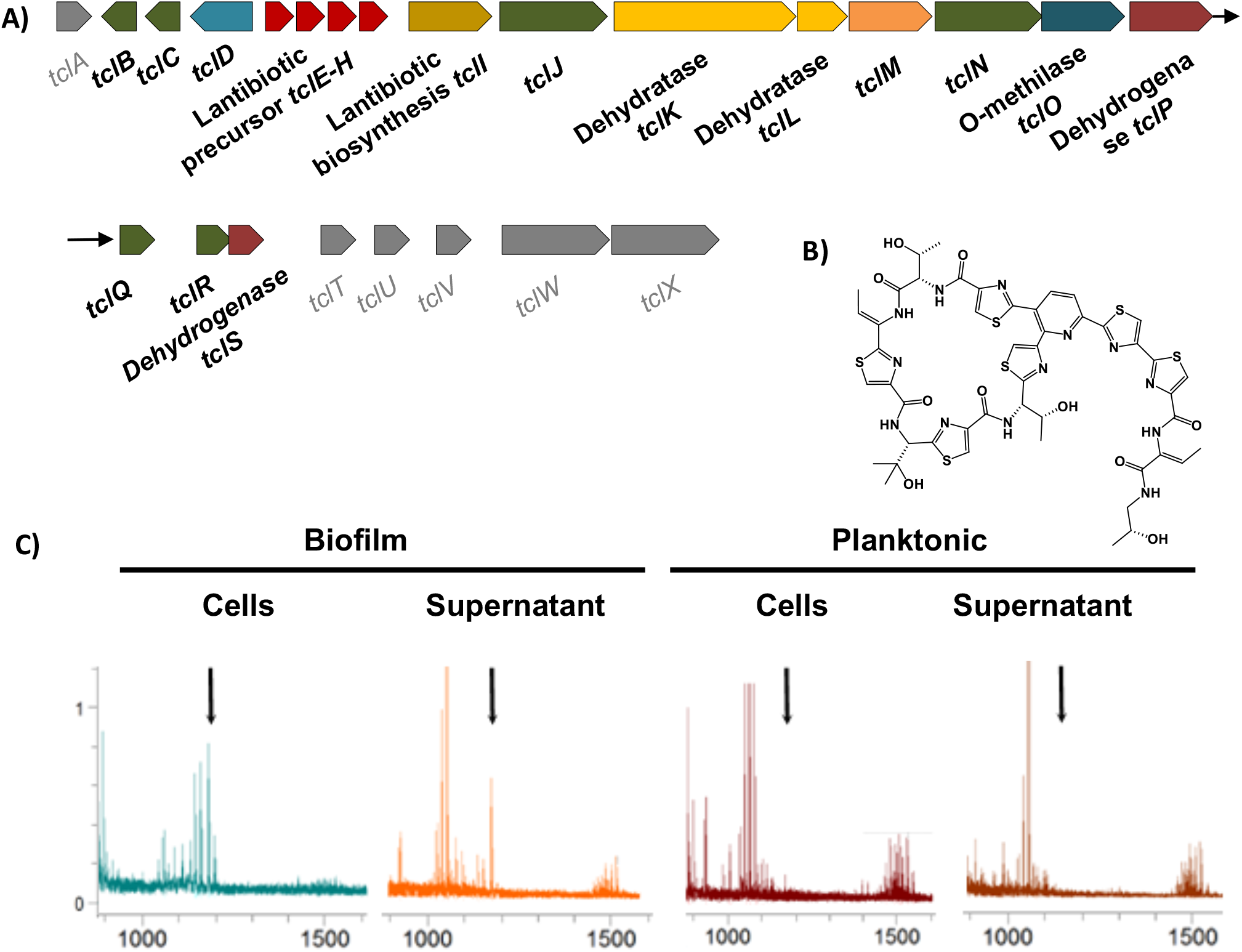
Cells in a biofilm produce more antimicrobials. **A)** Scheme of the genes for the biosynthesis of thiocillin are overexpressed in biofilm cells (*BC5094-BC5070*). Genes with no variation in expression level are shown in gray color. **B)** 3D structure of the thiocillin molecule. **C)** Detail of the traces from HPLC-MS-MS (TOF-TOF) of cells and supernatants of 48 h aged cultures. Biofilms were collected and suspended in PBS, separating cells from the supernatant after centrifugation. Culture medium was centrifuged to separate floating cells from the supernatant. Samples were purified with C8 ZipTip^®^ (Merck) previous to analysis. Black arrows indicate the positions of the thiocillin peaks. See the supplemental material for the complete spectra (Fig. S5).

Reactive oxygen species (ROS) are well-known factors which elicit antimicrobial-mediated killing and are discussed to be the final cause of bacterial cell death instead of the direct effect of antimicrobials [45]. ROS production has also been described to be part of the defensive response of plants and mammals to fight against pathogen invasions [46]. Bacteria produce several enzymes which neutralize ROS, either preventing or reducing cell damage as well as other repairing enzymes. According to the protective mode which characterizes the lifestyle of biofilm cells, our analyses revealed the overexpression of enzymes devoted to scavenging diverse ROS: catalase (BC3008), superoxide dismutases (*BC1468, BC4907*), chloroperoxidase (*BC4774*), alkylhydroperoxidase (*BC2830*,) and glyoxalases (*BC5092*, BC3178, BC0824*). Additionally, iTRAQ analysis showed higher amounts of glutathione peroxidase (*BC2114*), glyoxalases (*BC5092, BC3532*), superoxide dismutase (*BC4272*), thioredoxin-dependent thiol peroxidase (*BC0517*), thiol peroxidase (*BC4639*) and heme-dependent peroxidase (*BC5388)* in biofilm-associated cells (Fig. 5A). In this direction we additionally observed: i) downregulation of the TCA cycle (*BC1251*, BC3833*, BC3834*, BC1790*, BC0466*, BC1712*, BC0466\**), including Complex II of the electron transport chain (BC4516*, BC4517*), which supplies two electrons to ubiquinone (Fig. 5B and Fig. S6); ii) overexpression of Complex I of the electron transport chain, increasing the total amount of the flavin mononucleotide (FMN) which receives the electron from NADH [47, 48]; and iii) activation of the glyoxylate shunt (isocitrate lyase *BC1128\**, malate synthase *BC1127\**), a bypass of the TCA which converts isocitrate to glyoxylate to malate. This shunt yields less reducing power, and indirectly increases the concentration of NAD^+^, which have been proven to arrest superoxide production by Complex I, and complementary being implicated in the level of tolerance to antibiotics [49–53].

**Fig 5.**
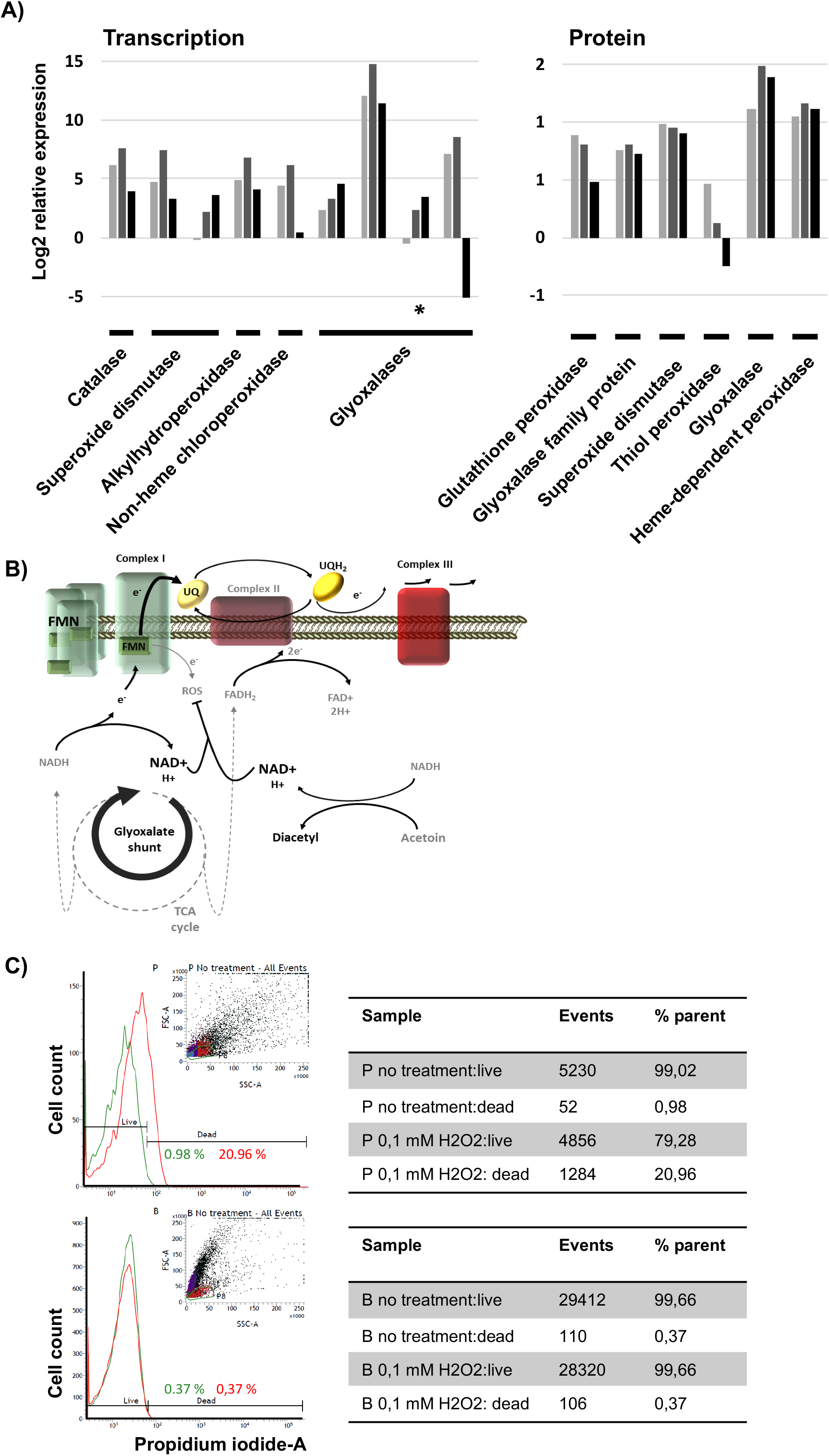
Cells in a biofilm trigger the machinery dedicated to scavenging reactive oxygen species. **A)** Expression pattern of ROS detoxification enzymes in biofilm compared to floating cells at 24 h (light gray), 48 h (gray) and 72 h (black) based on RNAseq (left) or iTRAQ (right) analysis. **B)** Scheme of the metabolic rearrangement to reduce ROS production from the electron leakage of the flavin mononucleotide (FMN). Overexpression of the complex I and FMN, reduction of the NADH/NAD^+^ ratio degrading acetoin, activation of the glyoxylate shunt, and reduction of the ubiquinone (UQ) saturation lead to release the fully reduced state of the FMN, preventing electron leakage. **C)** ROS survival assay of planktonic (top) and biofilm (bottom) cells of 48 h aged after treatment with 0.5 mM H_2_O_2_ solution for 30 minutes. Cell viability was measured by cell staining with Live and Dead^®^ and analysis with flow cytometry comparing each untreated population (green lines) with treated biofilm or planktonic cells (red lines).

To test the major resilience of biofilm cells than planktonic cells to damage induced to ROS stress, we evaluated the response of the two populations at 48 h aged to 0.1 mM H2O2 solution. After 30 minutes of exposure, planktonic cells showed around 20 % of mortality while in the biofilm population was unaffected (Fig. 5C), supporting the idea of the preparation of the internal machinery that sessile bacterial cells program to defend against putative damage mediated by ROS. A comparable cell numbers of each subpopulation was initially used for the treatment, however, we found a reduction in the number of event counts in the planktonic subpopulation in the flow cytometry analysis, a fact that might be altering the real effect of oxidative stress and should be addressed in future works.

### Planktonic cells express much higher levels of virulence factors

In the interaction with hosts, bacterial cells must overcome their diversified immune responses, and this is achievable with the gain of so-called virulence factors. We have found two factors which might complementarily contribute to the survival of bacterial cells despite the host immune system. One factor is a specific beta-lysine acetyltransferase (*BC2249\**), which may confer resistance to beta-lysine, an antibacterial compound produced by platelets during coagulation that induces lysis in many Gram-positive bacteria [54]. The other factor is the immune Inhibitor A (InhA) precursor, which is a bacterial enzyme able to i) digest attacins and cecropins, two classes of antibacterial humoral factors in insects [55], ii) cleave hemoglobin and serum albumin, two natural sources of amino acids which are thus relevant to bacteremia [56]; and iii) InhA1 is a key effector implicated in the liberation of spores from macrophages [57]. Consistent with the biofilm resilience and the enhancement of sporulation within the biofilm, the three InhA paralogues (*BC0666^x^, BC1284*, BC2984\**) found in the genome of *B. cereus* are upregulated in biofilm. The expression of InhA2 (*BC0666*) is dependent of PlcR, which is downregulated, what is contradictory. However, iTRAQ results indicated that *BC0666* quantity of protein was reduced, reinforcing the rationale of running additional and complementary analysis as iTRAQ to confirm the findings of RNAseq.

In addition to these changes in gene expression leading to resilience of *B. cereus* biofilms against the host immune system, *B. cereus* biofilm cells seems to reduce their toxicity to host. This bacterium possesses a variety of toxins: cytotoxin K (cytK), non-hemolytic enterotoxin (NHE), enterotoxin C (EntC), hemolysin III, hemolysin BL (HBL), haemolysin XhlA, invasins (three collagenases and a phospholipase C), and thiol-activated cytolysin I. These toxins and proteases and its positive regulator PlcR are strongly downregulated in biofilm-associated cells, except for hemolysin III, which is independent of PlcR, [58]. To demonstrate that floating cells are more prone than biofilm-associated cells to attack the host, both bacterial cell types were isolated and tested for their toxicity potential toward human HeLa and MDA cell lines in toxicity assays (Fig. 6 and supplemental movie). As expected from our analysis, floating cells, which possess an enhanced machinery for toxin production, killed human cells much faster than biofilm-associated cells did, an effect seen using different bacteria:cell ratios in both cell lines. These enterotoxins are labile and do not survive stomach passage, injuring the hosts only when they are produced within the intestine. Thus, it might be reasonably suggested that floating cells and germinating spores of *B. cereus* in the transit through the intestinal track are prone to produce the toxins; and biofilm cells are more oriented to defense and inhibition of the immune response. The strategy used by floating cells seems to be less oriented to survive the host attack, redirecting all the efforts to produce toxins and other virulence factors with the aim of achieving a fast, aggressive and effective attack to the host, as reported in some clinical cases in which patients died within several hours after the intake of contaminated food [59].

**Fig 6.**
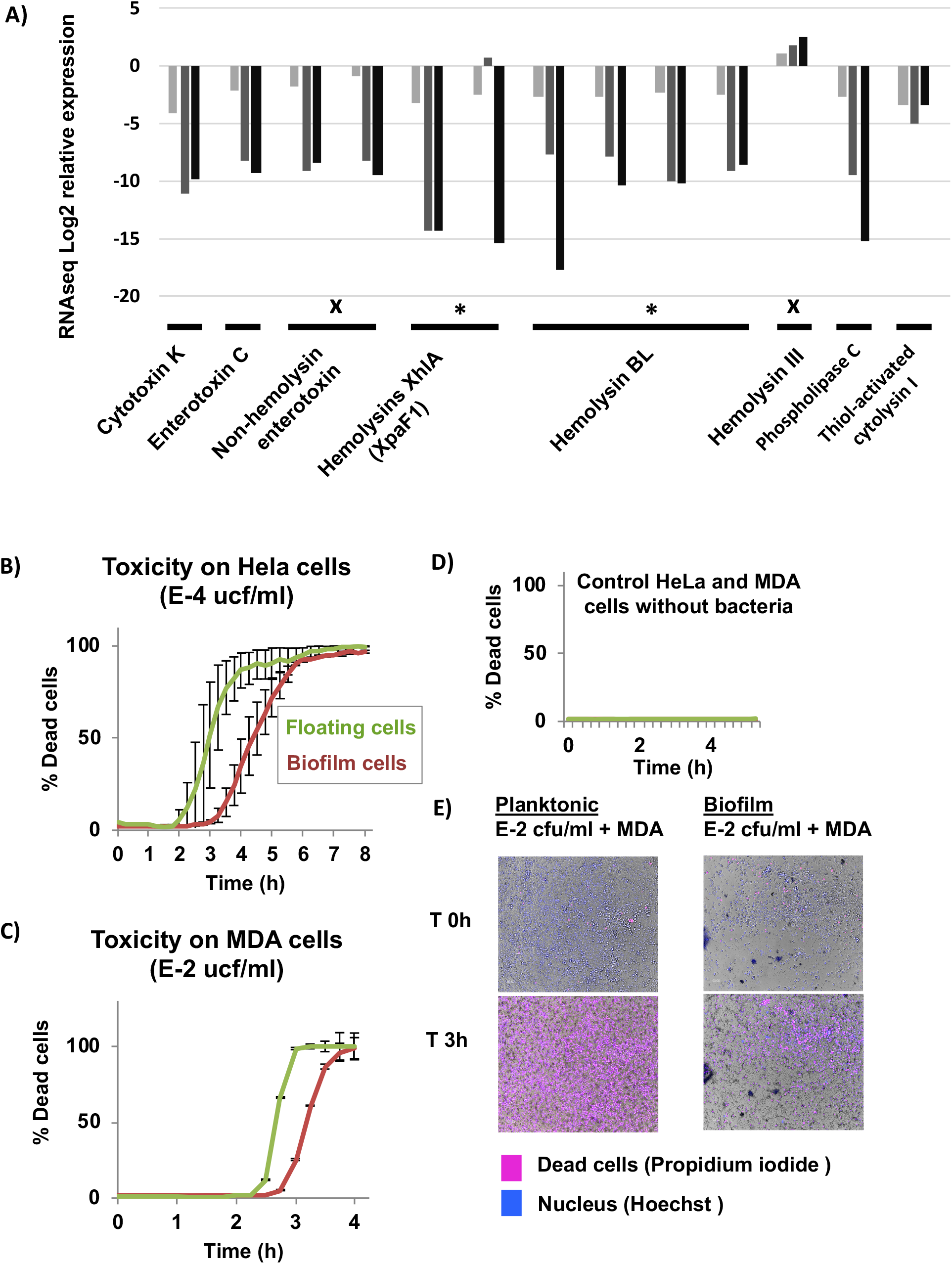
Cells in a biofilm are less aggressive to the host than floating cells are. **A).** Expression pattern of toxins in a biofilm compared to floating cells at 24 (light gray), 48 (gray) and 72 h (black) (RNAseq data). Data confirmed by iTRAQ results are marked with a star. Data in conflict with iTRAQ data are marked with a cross. **B-C)** Toxicity assay of floating (green line) or biofilm (red line) cells of 48 h cultures against HeLa and MDA cell lines culture without addition of antibiotics. Error bars indicate SD (n=12). **D)** Negative control of the toxicity assay. **E)** Micrographs of the toxicity assay after 3 h of incubation. Dead cells (pink) are stained with propidium iodide. Nuclei are stained with Hoechst (blue). (See the attached videos of cell death progress in the supplemental material).

### Planktonic and sessile cells coordinately attack and defend

In our experimental setup, biofilm and floating cells coexist, which led us to explore the existence of metabolic specialization and possible connections between these two populations of cells at 24 h, when each population initiates the differentiation process. Among the genes showing differential expression between floating and biofilm cells, 862 showed predicted associations in the STRING database with a confidence threshold > 500, which were used to build a primary network using the STRINGdb and iGraph R packages (Fig. 7). Using the Betweenness algorithm (Newman and Girvan), we found 16 subnetworks with a minimum of ten genes. We observed that the majority of modules contain genes either more highly expressed in the biofilm (red nodes) or in floating cells (blue nodes), corresponding to specific functions described previously, which clearly belong to biofilm or planktonic cells. Two examples of clusters supported the validity of our analysis, showing enrichment of expected functions for biofilm or floating cells, respectively: Cluster 2 contains genes associated with sporulation, a developmental process that is upregulated in biofilms (Cluster 2: Fig. 7 and Supplemental Table S6). Cluster 3 is involved in the bacterial flagella assembly and chemotaxis, a function that is predominantly activated in floating cells (Cluster 3: Fig. 1C and Supplemental Table S7). Clusters containing mixed elements (red for biofilm and blue for floating cells) were ranked by size and functionally analyzed to identify pathways that could be existent in both cell populations. The mixed Cluster 1 is composed of 83 genes (52 planktonic and 32 biofilm). A total of 9 KEGG pathways are included in this cluster (Supplemental Table S3). Within these global metabolic groups, the integration of genes from each population is specifically outstanding for: i) the TCA cycle and 2-oxocarboxylic acid metabolism, with the floating and biofilm genes clearly divided into different sub-routes within the same metabolic pathways (Fig. S7-S8) and ii) three metabolic KEGG pathways containing 85% of genes that have known functions (Cluster 1 Fig. 7 and Supplemental Table S4-S5) and that are prominently dedicated to secondary metabolism and antibiotic biosynthesis.

**Fig. 7.**
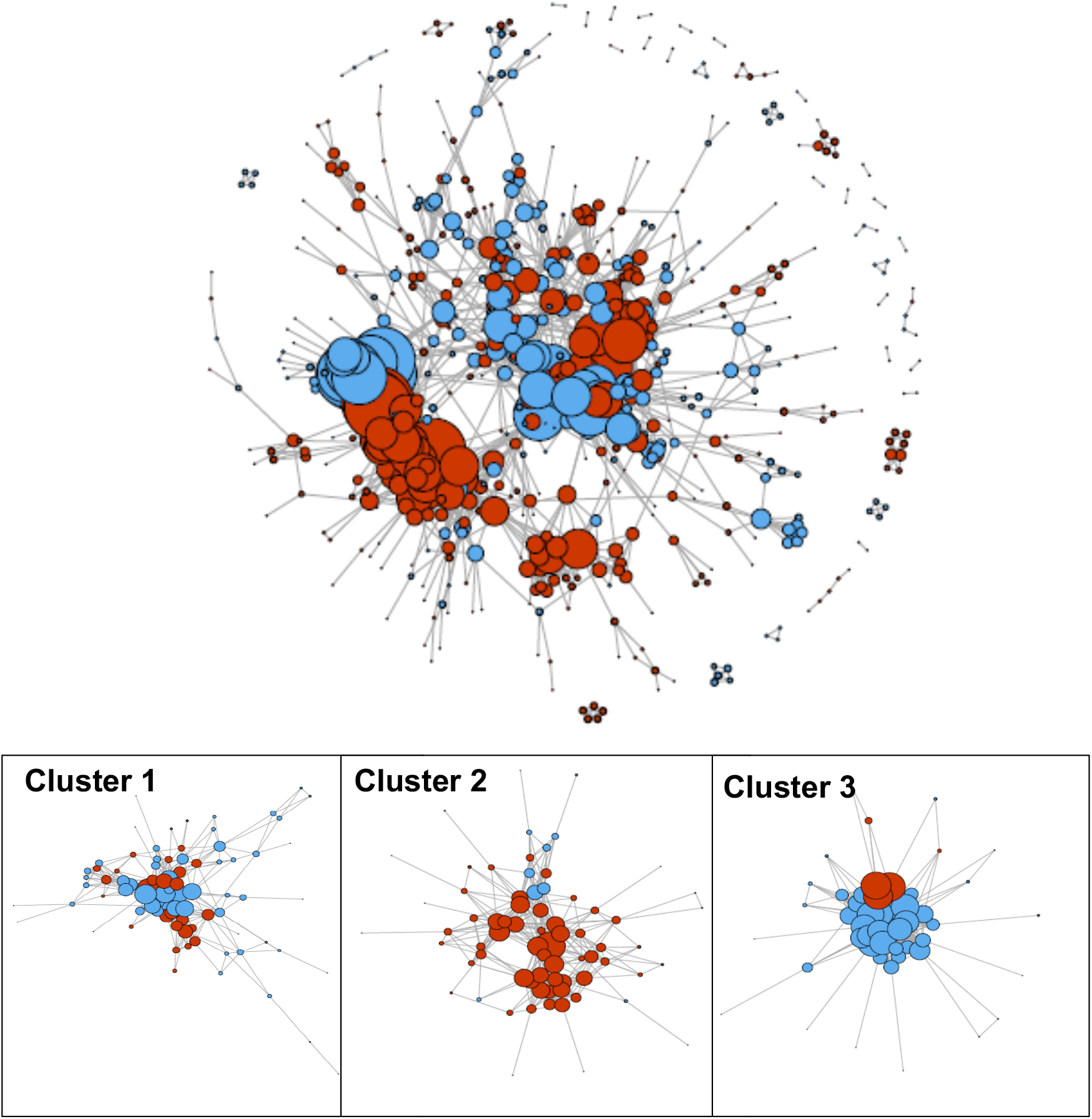
Specialization of biofilm and floating cells in distinct but coordinated metabolic activity. Networks formed by the differentially expressed (DE) genes and the functional associations between them. Each node represents a gene significantly DE between planktonic and biofilm samples at 24 hours; each edge represents an association between two genes according to the STRING database, as described in the Methods. Red nodes represent genes more highly expressed in the biofilm samples; blue genes represent genes more highly expressed in the planktonic samples. To the left is the global network formed by all DE genes. To the right, Clusters 1-3 represent subsets of the global network, obtained using the module detection method. Cluster 1 contains genes related to sporulation and are mostly from biofilm cells. Cluster 2 contains genes dedicated to flagellum assembly and chemotaxis and are mostly from floating cells. Cluster 3, contains a mix of genes from both populations (floating and biofilm cells) and are dedicated to global metabolic pathways.

All the information obtained from our study have been integrated into a model which collects the most distinctive features characterizing floating or biofilm-associated cells (Fig. 8). Biofilm-associated cells seem to direct the energy to the synthesis of the extracellular matrix and anatomical changes conducive to individual resilience to external aggressions. This specialization is complemented by the deviation of the metabolism to the synthesis of secondary metabolites, some of which might mediate the interaction with competitors at short distances from or even in close contact with the biofilm (Fig. 6B). However, floating cells are metabolically predisposed to colonize new niches and are also more aggressive in terms of pathogenicity, showing an increased production of toxins, conducive to the acquisition of more nutrients. In summary, this study redefines our view of *B. cereus* biofilm versus floating cells and goes further, mapping how deep the physiological and functional changes are in the switch from one lifestyle to the other.

**Fig 8.**
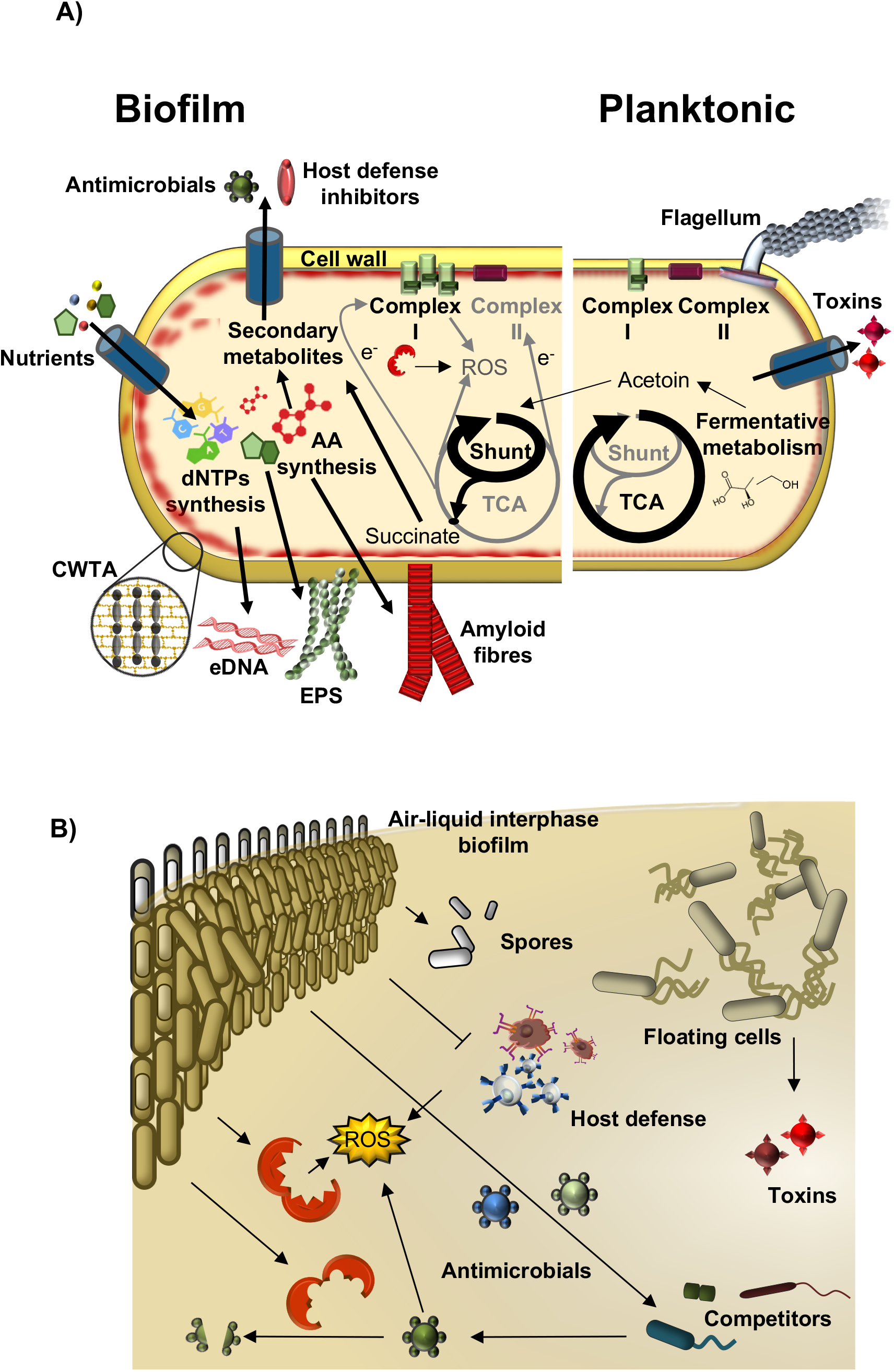
Model of offence and defense of floating cells and cells in the air-liquid interphase biofilm. **A)** Major changes differentiate the biofilm from floating cells. Biofilm cells manifest a collection of physiological and morphological changes leading to protection mediated by the increase in the cell wall thickness, the expression of inhibitors of the host defense and antimicrobial resistance genes, or the triggering of sporulation and enhancing of the ROS detoxification system. **B)** These changes coordinate with an offensive action against competitors driven by the production of antimicrobials. In contrast, floating cells tend to be more active metabolically and more aggressive in the interaction with hosts, overexpressing, among other compounds, a battery of toxins. CWTA, cell wall teichoic acids; EPS, exopolysaccharides.

## Methods

### Bacterial Strains and Culture Conditions

The bacteria used in this study was *B. cereus* ATCC14579 obtained from the Spanish collection of type strains (CECT148). The mediums used to culture bacteria were LB agar and liquid TyJ broth [(1% tryptone, OXOID), 0.5% yeast extract (OXOID), 0.5% NaCl, 10 mM MgSO4, and 0.1 mM MnSO4] were used for bacterial cultures. Bacteria were routinely streaked from −80°C stocks onto LB agar and incubated at 24 h at 30°C before each experiment. For biofilm experiments, one colony was suspended in 1 ml of TyJ broth, and the final OD600 was adjusted to one. One milliliter of TyJ in 24-well plates was inoculated with 10 μl of the bacterial cell suspension. Only central lines of the wells in the plates where used for culture. Plates were incubated at 30°C for 24, 48 and 72 h.

### Cell sampling, RNA isolation and whole transcriptome sequencing

Biofilm, a bacterial biomass adhered to the walls of the wells, was collected with sterile cotton swabs, suspended in 1 ml of TyJ, and centrifuged at 12000 g for 10 seconds; the pellet was immediately frozen in liquid nitrogen after discarding the supernatant. As the amount of biofilm formed change over time, collections from 2-8 wells were merged to reach a feasible number of cells for further RNA isolation. For floating cell sampling, 250-500 μl of culture was collected from several wells without disturbing the submerged biofilm, mixed in a 2 ml tube, and centrifuged at 12000 g for 10 seconds; the pellet was immediately frozen in liquid nitrogen after discarding the supernatant. Samples were stored at −80°C. Samples were recovered from −80°C, and cold beads were added to each tube and 900 μl of TRI-reagent (Invitrogen), followed by bead beating in a tissue lyser immediately for 1 min. The samples were placed 3 minutes at 55°C. After addition of 200 μl chloroform, tubes were vortexed for 10 seconds, incubated 2-3 min at room temperature and centrifuged at 12,000 g for 10 minutes at 4°C. The upper phase was mixed with 500 μl ice-cold isopropanol for RNA precipitation, incubated for 10 minutes at RT, and centrifuged at 12,000 g for 10 minutes at 4°C. The pellet was washed with 75% ethanol, centrifuged twice to remove all supernatant and air-dried for 5 minutes. The pellet was suspended in 20 μl DEPC water (Carl Roth GmbH). DNA was removed using RQ1 DNase treatment (Promega) with a RiboLock RNase inhibitor (Thermo) following the product instructions. After digestion, 400 μl DEPC-MQ water, 250 μl phenol and 250 μl chloroform were added to the samples, vortexed for 5 seconds and centrifuged at 15000 g for 15 minutes at 4°C. The supernatant was mixed with 1 ml ice-cold 10% ethanol and 50 μl 3 M sodium acetate pH 5.2. The mixture was incubated at least 30 minutes at −20°C and then centrifuged at 15000 g for 15 minutes at 4°C. The pellets were washed twice with 75% ethanol and centrifuged at 15000 g for 5 minutes at 4°C. After spinning the tubes to retire as much of the supernatant as possible, pellets were air-dried at RT for 5 minutes and suspended in 20 μl DEPC water (Carl Roth GmbH). Samples were sent as two biological replicates to the PrimBio Research Institute (Exton, PA, USA), at which the 16S/23S rRNA removal with the Ribo-Zero kit, mRNA quality control (QC), cDNA library preparation, library QC, template preparation, template QC, and RNA-sequencing on an Illumina platform was performed.

### Data analysis of the whole transcriptomes

After trimming the raw RNA-seq reads (PE150 (paired-end 150 bp)) from the adapter sequences, reads were mapped against the *B. cereus* ATCC 14579 genome sequence. The RPKM (reads per kilo base per million mapped reads) table was generated, and differential gene expression analyses were carried out with the webserver pipeline T-Rex [60] on the Genome2D webserver (http://genome2d.molgenrug.nl/). The significance threshold was defined by a p-value□of <□0.05 and a fold-change of >2 (“TopHits” in T-REx). Cluster of orthologous groups (COG) analysis was performed using the Functional Analysis Tools of the T-REx pipeline. The RNA-seq data from this study have been submitted to the NCBI Gene Expression Omnibus (GEO; http://www.ncbi.nlm.nih.gov/geo/).

BAGEL3 (reference) and antiSMASH (reference) were used to identify secondary metabolite synthesis regions in the genome of *B. cereus* ATCC 14579. To compare the behavior of sporulation genes across the maturation progress of the biofilm, we used *k*-means clustering with the STEM (Short Time-series Expression Miner) software package.

### Proteomic analysis

Biofilm and floating cells were separated following the method described above. Samples were sent in triplicate to the Proteomic Facility of the Centro Nacional de Biotecnología CSIC (Spain) for Isobaric Tags for Relative and Absolute Quantitation (iTRAQ). Protein digestion and tagging with a TMTsixplex™ reagent was performed as follows: The total protein concentration was determined using a Pierce 660 nm protein assay (Thermo). For digestion, 40 μg of protein from each condition was precipitated by the methanol/chloroform method. Protein pellets were suspended and denatured in 20 μl 7 M urea/2 M thiourea/100 mM TEAB, pH 7.5, reduced with 2 μL of 50 mM Tris(2-carboxyethyl) phosphine (TCEP, SCIEX), pH 8.0, at 37°C for 60 min and followed by 1 μL of 200 mM cysteine-blocking reagent (methyl methanethiosulfonate (MMTS, Pierce) for 10 min at room temperature. Samples were diluted up to 140 μL to reduce the urea concentration with 25 mM TEAB. Digestions were initiated by adding 2 μg of sequence grade-modified trypsin (Sigma-Aldrich) to each sample at a ratio of 1:20 (w/w); the samples were then incubated at 37°C overnight on a shaker. Sample digestions were evaporated to dryness in a vacuum concentrator. The resulting peptides were subsequently labeled using a TMT-sixplex Isobaric Mass Tagging Kit (Thermo Scientific, Rockford, IL, USA) according to the manufacturer’s instructions, as follows: 126: 1B-24 h; 127: 1B-48 h; 128: 1B-72 h; 129: 1P-24 h; 130: 11B-24 h; 131: 13B-72 h. After labeling, the samples were pooled, evaporated to dryness and stored at −20°C until the LC-MS analysis.

For liquid chromatography and mass spectrometry analysis, 1 μg of the labeled protein mixture was subjected to 1D-nano LC ESI-MS/MS analysis using a nanoliquid chromatography system (Eksigent Technologies NanoLC Ultra 1D plus, SCIEX, Foster City, CA) coupled to a high-speed Triple TOF 5600 mass spectrometer (SCIEX, Foster City, CA) with a Nanospray III source. The analytical column used was a silica-based reversed-phase ACQUITY UPLC□ M-Class Peptide BEH C18 Column, 75 μm × 150 mm, 1.7 μm particle size and 130 Å pore size (Waters). The trap column was C18 Acclaim PepMap™ 100 (Thermo Scientific), 100 μm × 2 cm, 5 μm particle diameter, 100 Å pore size, switched on-line with the analytical column. The loading pump delivered a solution of 0.1% formic acid in water at 2 μl/min. The nanopump provided a flow-rate of 250 nl/min and was operated under gradient elution conditions. Peptides were separated using a gradient of 250 minutes ranging from 2% to 90% mobile phase B (mobile phase A: 2% acetonitrile, 0.1% formic acid; mobile phase B: 100% acetonitrile, 0.1% formic acid). The injection volume was 5 μl.

Data acquisition was performed with a TripleTOF 5600 System (SCIEX, Foster City, CA). Data were acquired using an ionspray voltage floating (ISVF) 2300 V, curtain gas (CUR) 35, interface heater temperature (IHT) 150, ion source gas 1 (GS1) 25, declustering potential (DP) 150 V. All data was acquired using information-dependent acquisition (IDA) mode with Analyst TF 1.7 software (SCIEX, Foster City, CA). For IDA parameters, a 0.25 s MS survey scan in the mass range of 350–1250 Da were followed by 30 MS/MS scans of 150 ms in the mass range of 100–1800. Switching criteria were set to ions greater than a mass-to-charge ratio (m/z) of 350 and smaller than m/z 1250 with a charge state of 2–5 and an abundance threshold of more than 90 counts (cps). Former target ions were excluded for 20 s. An IDA rolling collision energy (CE) parameters script was used for automatically controlling the CE.

For data analysis and protein identification, the mass spectrometry data obtained were processed using PeakView^®^ 2.2 software (SCIEX, Foster City, CA). Raw data file conversion tools generated mgf files, which were also searched against the *B. cereus* protein database from Uniprot (database state June-August 2016), containing 40530 protein-coding genes that included their corresponding reversed entries using four different search engines (Mascot, OMSSA, X!TANDEM and MyriMatch). Search parameters were set as follows: enzyme, trypsin; allowed missed cleavages, 2; methylthio (C) as a fixed modification and TMT-6plex (N-term, K, Y), acetyl (protein N-term), oxidation (M), Gln->pyro-Glu (N-term Q) and Glu->pyro-Glu (N-term E) as variable modifications. The peptide mass tolerance was set to ± 25 ppm for precursors and 0.02 Da for fragment masses. The confidence interval for protein identification was set to ≥ 95% (p<0.05), and only peptides with an individual ion score above the 1% false discovery rates (FDR) threshold were considered correctly identified. A 5% quantitation FDR threshold was estimated to consider the significant differentially expressed proteins.

### Network creation, clustering and functional analysis

Data on potential pairwise associations between *Bacillus cereus* genes were downloaded from the STRING database using the STRINGdb Bioconductor package for R. This resource comprises data from multiple sources related to direct interactions, both direct and inferred through homology, and predicted associations, made through a variety of algorithms using different data sources. These pairwise associations were then used to build a network between all genes that showed differential expression between biofilm and floating samples, using iGraph, visualized using Cytoscape. This network was then clustered using the edge betweenness algorithm (Newman and M Girvan, 2004) to find modules of genes with a high degree of connectivity between them, compared to connections to genes outside the module. The DAVID tool was employed for the genes’ functional annotation and to look for enrichment of specific biological processes in the GO and KEGG databases within the cluster. When GO and KEGG enrichment analyses showed the same pathways, the results of one of these two database sources were selected to avoid functional annotation redundancy of clusters.

### Mass spectrometry analysis for thiocillin detection

Thiocillin detection was analyzed using HPLC-MS-MS (Ultraflex TOF-TOF, Bruker) of cells and supernatants of 48 h cultures of *B. cereus* in TyJ medium incubated at 28°C without shaking. The biofilm was collected and thoroughly suspended in PBS and then centrifuged at 12000 g to separate cells from the supernatant. Culture medium was centrifuged to separate floating cells from the supernatant. Previous to analysis, the samples were purified with C8 ZipTip^®^ (Merck) to discard salts. To perform MS-MS, a low molecular weight matrix was used.

### Construction of mutants in *B. cereus*

*B. cereus eps1* or *eps2* mutant were obtained by electroporation using the plasmid pMAD, harboring a fragment to delete the genes *BC5279-BC5274* or genes *BC1583-BC1591* respectively by double recombination. The construct was created by joining PCR. In the first step, regions flanking the target genes were amplified separately, purified, and used for the joining PCRs. These PCR products were digested and cloned into the pMAD vector digested with the same enzymes. The resulting suicide plasmids were used to transform *B. cereus* electrocompetent cells as described previously with some modifications.

Complementation of *eps2* mutant was done using the replicative plasmid pUTE973 (kindly provided by Theresa M. Koehler, University of Texas) harboring the construction of *P_IPTG_-eps2*. The primers used to amplify the genomic region were *eps2.Fw TTTTGTCGACGATGAAGAGATACGAGGAATTGG* and *eps2.Rv TTTTGCATGCGTTGAAACCAAATTACAATCTC* and the fragment was cloned in the SalI and SphI cut of pUTE973 downstream of the *P_iptg_* promoter. The activity of this promoter was triggered using a 1mM of IPTG solution.

Electroporation was performed with 10 μg of plasmids in 100 μL of electrocompetent *B. cereus* in 0.2-cm cuvettes using the following electroporation parameters: voltage 1400 kV, capacitance 25μF, resistance 400 Ω. After electroporation, cells were incubated in LB for 5 hours, and then seeded in LB medium supplemented with X-Gal and erythromycin for 72 h at 30°C. Blue colonies were selected and streaked to trigger allele replacement. Finally, white colonies that were sensitive to MLS were selected, and deletion of the target gene was verified by colony PCR analysis and sequencing of the amplicons.

### Evaluation of bacterial cell wall thickness

Biofilm and floating cells were separated following the method described above and fixed in 2% glutaraldehyde. Post fixation was performed with 1% osmium tetroxide in 0.1 M, pH 7.4 phosphate buffer, following dehydration in an acetone gradient at 4 °C: 30%, 50%, 70% and 90%. A step for in bloc staining with 1% uranyl acetate in 50% cold acetone was included after the 50% step, leaving the samples overnight at 4 °C. Dehydration was continued with serial incubations in absolute acetone and propylene oxide at room temperature. The embedding in Spurr’s resin was made following different steps that combined Spurr’s resin:propylene oxide at 1:1, 3:1 (overnight) and two changes in pure resin (the second one, overnight). Finally, samples were embedded in pure resin at 70 °C for 3 days. Ultrathin sections were visualized in a JEOL JEM-1400 transmission electron microscope with a high-resolution camera (Gatan ES1000 W). The analysis was done using ImageJ-Fiji software over 40 images of each sample and 6-10 measurements over each picture using only circular cell sections to avoid the effect of a tilted sectioning.

### ROS survival assay

Biofilm and planktonic populations were collected in NaCl buffer (0.5 g/L), mild sonicated for 30 seconds at 30% intensity to separate cells and exposed to 0.1 mM of H2O2 for 30 minutes. H2O2 was retired by centrifugation and cells were suspended again in NaCl buffer. Before cytometry analysis (BD FACSVerse^®^ equipment), cells received a sonication bath for 3 minutes. Cell deadness was assayed using Live and Dead^®^ reactive (Life Sciences) following the manufacturer instructions. The analysis was done over a population gated considering planktonic sample, which contains single cells, avoiding biofilm aggregates. Untreated biofilm and planktonic cells were used as controls. ROS survival was assayed by comparing the increase in mortality between treated bacteria with its own population without treatment.

### Human cell toxicity assay

MDA-MB-231 breast adenocarcinoma and HeLa cervical cancer cell lines were obtained from the American Type Culture Collection (ATCC) and were grown in RPMI 1640 and DEMEM glucose (4.5 g/L) medium cultures respectively, supplemented with glutamine (2 mM), penicillin (50 IU/mL), streptomycin (50 mg/L), amphotericin (1.25 mg/L), and 10% FBS, at 37°C with 5% CO_2_ in air. Cells were seeded at 2000 MDA and 1500 HeLa cells/well in a 96-well plate and incubated for 72 h at 37°C and 5% CO_2_ to achieve confluence and a cell density of 1·10^4^ cells/well. HeLa and MDA cell medium culture was replaced with ‘assay culture’ (supplemented with glutamine and FBS, without antibiotics), the cells were incubated for two hours, and then the culture medium was replaced again with assay culture.

*B. cereus* ATCC14579 was streaked onto an LB agar plate and incubated for 24 h at 28 °C. The *B. cereus* biomass was suspended in LB and seeded into a 24-well plate with 1 ml of TyJ and incubated for 48 h. Biofilm and floating cells were separated following the method described above. Both cell fractions of *B. cereus* were washed twice with sterile PBS, and the OD600 was adjusted to 1 (approx. 10^7^ cfu/ml). These bacterial fractions were serially diluted 2-10 times, inoculated into 96-well cell culture plates and centrifuged 5 minutes at 2000 g to force the bacteria to contact with human cell cultures. Propidium iodide (PI; 3μM final concentration) and DAPI (2.5 μM final concentration) were added to wells to check the viability state of the eukaryotic cells. The time-lapse assay was performed using an Operetta high content screening microscope (Perkin Elmer) using a 10x dry objective, maintaining plates at 37C and 5% CO_2_ throughout the assay. Fluorescence images for DAPI and PI were taken from each well every 15 minutes for 9+ hours. Cell death measurements were obtained from 3 well replicas/plate for each bacteria/cell combination. The percentage of dead cells in each well was counted automatically using Harmony software (Perkin Elmer). Mammalian nuclei were segmented based on their larger size (>60 um2) and higher DAPI staining intensities. Stuck together nuclei and the large bacterial aggregates were then filtered out by only selecting nuclei with a high degree of roundness (within 80% of a perfect circle). This segmented population was then analyzed for nuclear PI staining intensity, with only those nuclei higher than the PI threshold value (> 200) recorded as dead cells with the remainder scored as live cells.

### Crystal violet assay

*B. cereus* was grown in 1 ml of TyJ medium in 4.5 mm diameter plates. Biofilms were stained by removing the spent medium and filling the well with 2 ml of 1% crystal violet for 5 minutes, followed by three washing steps with deionized water. The crystal violet was then resuspended with 50% acetic acid. Finally, the absorbance was measured at 595 nm [16]. Statistical significance was assessed by repeated measures ANOVA with post-hoc paired Student’s t test. Results with p□0.05 were considered statistically significant.

## Ethics approval and consent to participate

Not applicable

## Consent to publish

Not applicable

## Availability of data and materials

Transcriptom ic data has been submitted to NCBI GEO: https://www.ncbi.nlm.nih.gov/geo/query/acc.cgi?acc=GSE115528

The mass spectrometry proteomics data have been deposited to the ProteomeXchange Consortium via the PRIDE [1] partner repository with the dataset identifier PXD010211

## List of abbreviations

RNAseq: RNA sequencing
iTRAQ: Isobaric tags for relative and absolute quantitation
COG: Cluster of Orthologous Groups
STEM: Short Time-series Expression Miner
eDNA: Extracellular DNA
EPS: Exopolysaccharide
TCA: tricarboxylic acid
LTA: Lipoteichoic acid
PG: Peptidoglycan
ROS: Reactive oxygen species
FMN: Flavin mononucleotide

## Author contributions

Conception or design of the work; DR, JCA

Acquisition, analysis, or interpretation of data for the work: DR, JCA, OK, EF, JAR, JRP

Drafting the work: DR, JCA

Revising it critically for important intellectual content: DR, JCA, OK, EF, AdV

Final approval of the version to be published: DR, JCA, OK, EF, JRP, AdV, JAR

## Acknowledgements

We would like to thank Anne de Jong (University of Groningen) for technical support in the use of T-Rex platform, and Ana María Álvarez (University of Málaga) for helping in biofilm experiments. We also thank members of the electron microscopy, cell biology and cell cultures, confocal microscopy (Dr. John Pearson, Bionand) and mass spectrometry services of SCAI (University of Málaga) and the Mass Spectrometry service of CNB (Centro Nacional de Biotecnología - CSIC, Spain) for technical support and guidance, and to Miguel Ángel Medina Torres for providing us with human cell lines. Joaquín Caro-Astorga is the recipient of a FPI contract [BES-2013-064134] from Ministerio de Economía y Competitividad. James Perkins is funded by Sara Borrell program from Carlos III (CD14/00242). Juan Antonio Ranea is funded by grants from Spanish Ministry of Economy and Competitiveness [SAF2016-78041-C2-1-R] and Andalusian Government with European Regional Development Fund [CTS-486]. The CIBERER is an initiative from the Instituto de Salud Carlos III. This work was supported by grants [AGL2012-31968 and AGL2016-78662-R] from Ministerio de Economía y Competitividad, Spanish Government and European Research Council Starting Grant under Grant [BacBio 637971].

