## Supplemental figures for "Biofilm formation displays intrinsic offensive and defensive features of *Bacillus cereus*"

### Slide 1
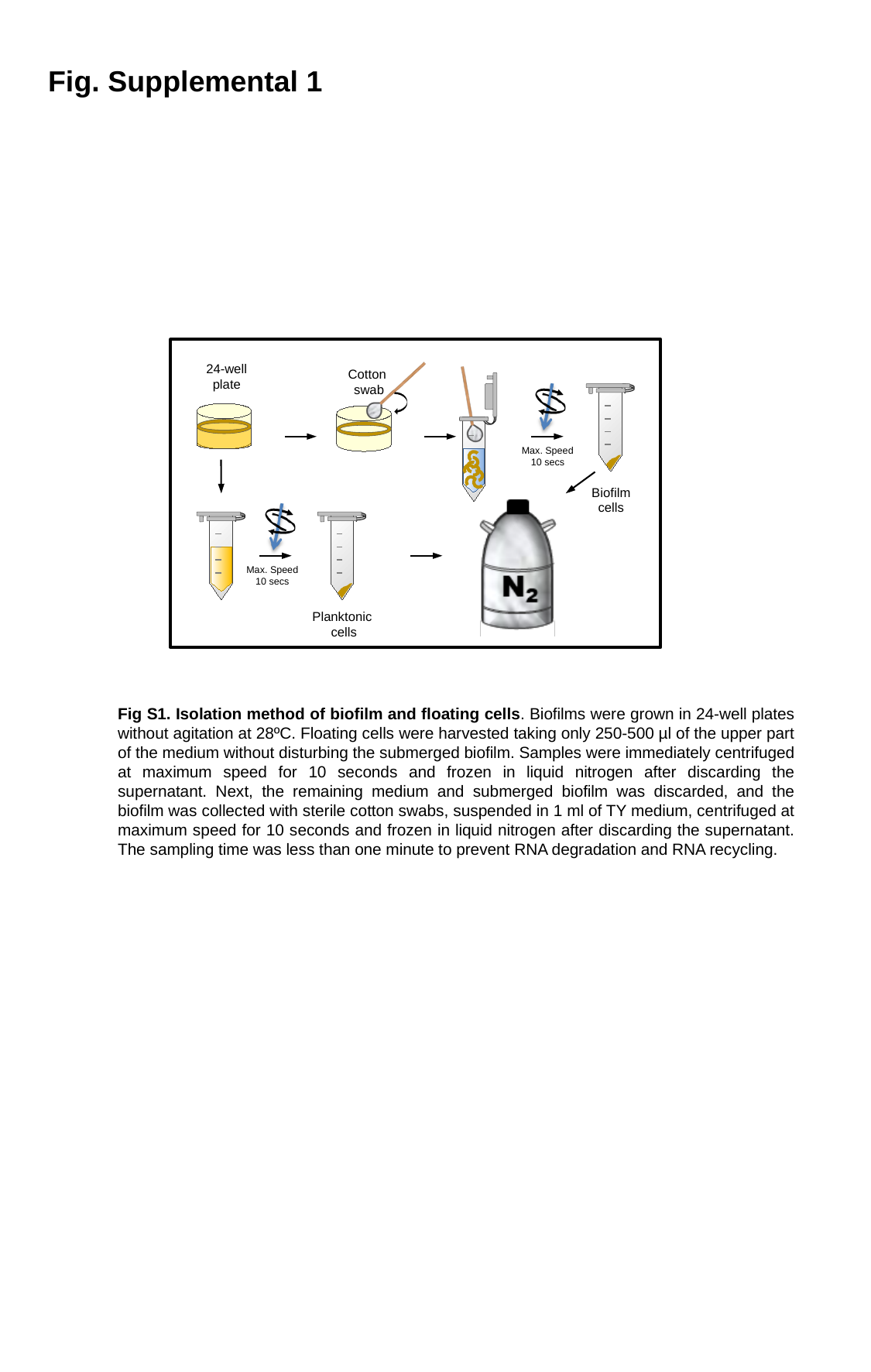

Fig. Supplemental 1
24-well plate
Cotton
swab
Max. Speed
10 secs
Biofilm
cells
Max. Speed
10 secs
Planktonic
cells
Fig S1. Isolation method of biofilm and floating cells. Biofilms were grown in 24-well plates without agitation at 28ºC. Floating cells were harvested taking only 250-500 µl of the upper part of the medium without disturbing the submerged biofilm. Samples were immediately centrifuged at maximum speed for 10 seconds and frozen in liquid nitrogen after discarding the supernatant. Next, the remaining medium and submerged biofilm was discarded, and the biofilm was collected with sterile cotton swabs, suspended in 1 ml of TY medium, centrifuged at maximum speed for 10 seconds and frozen in liquid nitrogen after discarding the supernatant. The sampling time was less than one minute to prevent RNA degradation and RNA recycling.

### Slide 2
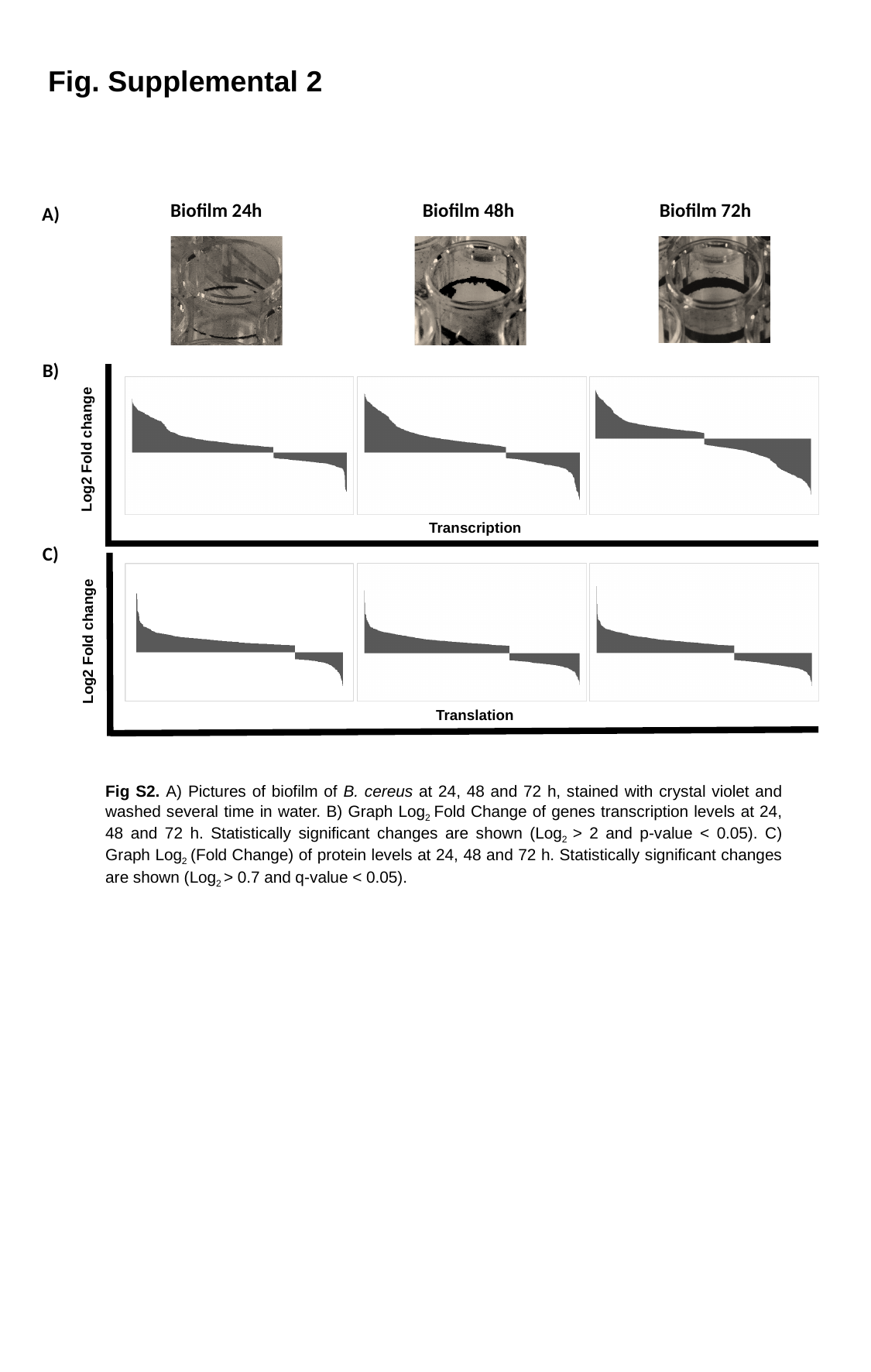

Fig. Supplemental 2
Biofilm 24h
Biofilm 48h
Biofilm 72h
A)
B)
Log2 Fold change
Transcription
C)
Log2 Fold change
Translation
Fig S2. A) Pictures of biofilm of B. cereus at 24, 48 and 72 h, stained with crystal violet and washed several time in water. B) Graph Log_2_ Fold Change of genes transcription levels at 24, 48 and 72 h. Statistically significant changes are shown (Log_2_ > 2 and p-value < 0.05). C) Graph Log_2_ (Fold Change) of protein levels at 24, 48 and 72 h. Statistically significant changes are shown (Log_2_ > 0.7 and q-value < 0.05).

### Slide 3
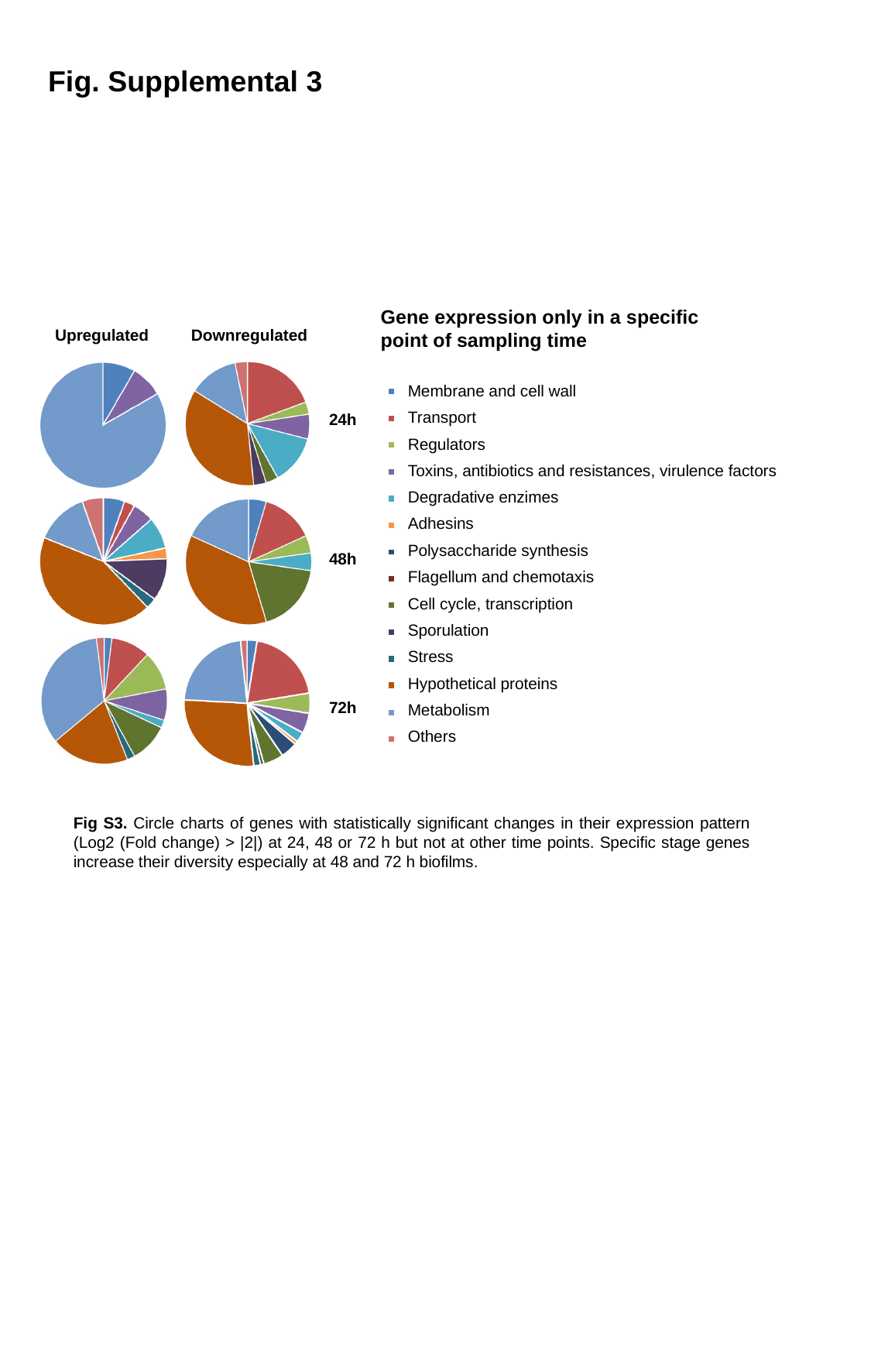

Fig. Supplemental 3
Gene expression only in a specific point of sampling time
Upregulated
Downregulated
Membrane and cell wall
Transport
Regulators
Toxins, antibiotics and resistances, virulence factors
Degradative enzimes
Adhesins
Polysaccharide synthesis
Flagellum and chemotaxis
Cell cycle, transcription
Sporulation
Stress
Hypothetical proteins
Metabolism
Others
24h
48h
72h
Fig S3. Circle charts of genes with statistically significant changes in their expression pattern (Log2 (Fold change) > |2|) at 24, 48 or 72 h but not at other time points. Specific stage genes increase their diversity especially at 48 and 72 h biofilms.

### Slide 4
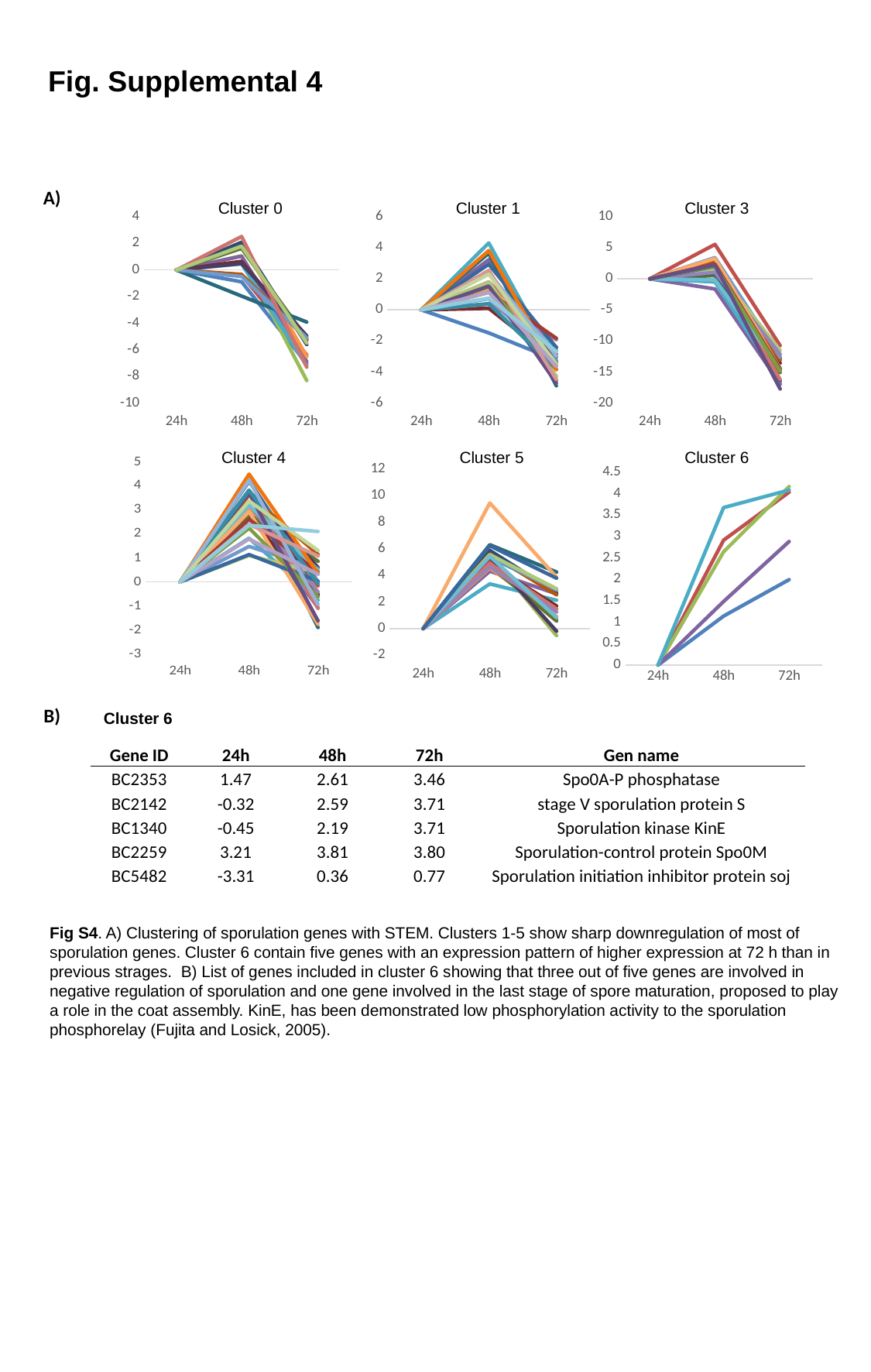

Fig. Supplemental 4
A)
Cluster 0
Cluster 1
Cluster 3
#### Chart
| Category | | | | | | | | | | | | | | | |
|---|---|---|---|---|---|---|---|---|---|---|---|---|---|---|---|
| 24h | 0.0 | 0.0 | 0.0 | 0.0 | 0.0 | 0.0 | 0.0 | 0.0 | 0.0 | 0.0 | 0.0 | 0.0 | 0.0 | 0.0 | 0.0 |
| 48h | -0.89 | -0.37 | 1.06 | 1.04 | 0.46 | 2.09 | 2.08 | 0.65 | 1.63 | 0.49 | -1.96 | -0.33 | -0.49 | 2.52 | 1.79 |
| 72h | -7.05 | -6.39 | -8.32 | -6.9 | -7.22 | -6.52 | -5.61 | -5.29 | -5.15 | -4.96 | -3.92 | -5.18 | -5.1 | -7.31 | -5.49 |
#### Chart
| Category | | | | | | | | | | | | | | | | | | | | | | | | | | | | | |
|---|---|---|---|---|---|---|---|---|---|---|---|---|---|---|---|---|---|---|---|---|---|---|---|---|---|---|---|---|---|
| 24h | 0.0 | 0.0 | 0.0 | 0.0 | 0.0 | 0.0 | 0.0 | 0.0 | 0.0 | 0.0 | 0.0 | 0.0 | 0.0 | 0.0 | 0.0 | 0.0 | 0.0 | 0.0 | 0.0 | 0.0 | 0.0 | 0.0 | 0.0 | 0.0 | 0.0 | 0.0 | 0.0 | 0.0 | 0.0 |
| 48h | -1.49 | 1.3 | 1.6 | 3.23 | 4.31 | 1.34 | 1.52 | 0.1 | 2.53 | 1.38 | 3.65 | 1.52 | 0.72 | 1.8 | 1.78 | 0.37 | 1.28 | 1.29 | 2.94 | 1.13 | 1.49 | 1.53 | 0.4 | 3.82 | 2.4 | 2.53 | 2.3 | 1.13 | 0.76 |
| 72h | -3.25 | -4.51 | -3.61 | -2.86 | -3.57 | -3.58 | -2.41 | -3.31 | -4.55 | -3.03 | -4.9 | -3.37 | -3.3 | -2.65 | -4.28 | -1.93 | -2.56 | -1.84 | -2.43 | -1.84 | -3.44 | -4.72 | -3.83 | -3.86 | -2.98 | -4.48 | -3.51 | -3.64 | -2.68 |
#### Chart
| Category | | | | | | | | | | | | | | | | | | | | | | |
|---|---|---|---|---|---|---|---|---|---|---|---|---|---|---|---|---|---|---|---|---|---|---|
| 24h | 0.0 | 0.0 | 0.0 | 0.0 | 0.0 | 0.0 | 0.0 | 0.0 | 0.0 | 0.0 | 0.0 | 0.0 | 0.0 | 0.0 | 0.0 | 0.0 | 0.0 | 0.0 | 0.0 | 0.0 | 0.0 | 0.0 |
| 48h | 2.65 | 5.55 | 1.57 | -1.64 | -0.5 | 2.38 | 3.03 | 2.75 | 0.53 | 2.5 | 0.85 | 2.9 | 3.42 | 1.91 | 1.67 | 1.05 | -0.17 | 3.28 | 2.0 | 2.35 | 2.1 | 2.24 |
| 72h | -16.99 | -10.77 | -14.54 | -16.92 | -16.49 | -12.89 | -12.68 | -13.58 | -15.11 | -16.39 | -14.41 | -13.07 | -12.62 | -16.24 | -11.52 | -12.16 | -14.35 | -14.48 | -14.58 | -14.59 | -14.86 | -17.76 |Cluster 4
Cluster 5
Cluster 6
#### Chart
| Category | | | | | | | | | | | | | | | | | | | | | | | | | | | | | |
|---|---|---|---|---|---|---|---|---|---|---|---|---|---|---|---|---|---|---|---|---|---|---|---|---|---|---|---|---|---|
| 24h | 0.0 | 0.0 | 0.0 | 0.0 | 0.0 | 0.0 | 0.0 | 0.0 | 0.0 | 0.0 | 0.0 | 0.0 | 0.0 | 0.0 | 0.0 | 0.0 | 0.0 | 0.0 | 0.0 | 0.0 | 0.0 | 0.0 | 0.0 | 0.0 | 0.0 | 0.0 | 0.0 | 0.0 | 0.0 |
| 48h | 4.19 | 2.87 | 3.72 | 2.84 | 1.82 | 3.08 | 4.25 | 2.68 | 2.8 | 3.79 | 3.82 | 3.62 | 1.48 | 3.71 | 1.1 | 3.22 | 3.2 | 2.93 | 1.14 | 2.61 | 2.24 | 3.7 | 3.81 | 4.49 | 4.24 | 2.4 | 3.37 | 1.79 | 2.36 |
| 72h | 0.01 | -0.16 | -0.43 | -1.07 | -0.76 | -0.01 | -0.57 | -0.52 | 0.86 | 0.6 | -1.9 | 1.15 | 0.54 | -1.1 | 0.38 | -0.42 | 0.5 | -1.76 | 0.03 | 0.4 | -0.62 | -1.62 | -0.07 | 0.36 | -0.93 | 1.08 | 1.31 | 0.32 | 2.1 |
#### Chart
| Category | | | | | | | | | | | | | | | | | | | |
|---|---|---|---|---|---|---|---|---|---|---|---|---|---|---|---|---|---|---|---|
| 24h | 0.0 | 0.0 | 0.0 | 0.0 | 0.0 | 0.0 | 0.0 | 0.0 | 0.0 | 0.0 | 0.0 | 0.0 | 0.0 | 0.0 | 0.0 | 0.0 | 0.0 | 0.0 | 0.0 |
| 48h | 5.37 | 5.06 | 5.51 | 4.32 | 3.36 | 4.56 | 5.88 | 4.85 | 4.73 | 5.4 | 6.31 | 5.62 | 5.44 | 4.9 | 5.56 | 4.64 | 5.44 | 9.45 | 6.23 |
| 72h | 2.93 | 0.73 | -0.53 | 2.61 | 2.13 | 0.9 | 2.69 | 1.7 | 0.57 | -0.2 | 4.25 | 2.54 | 2.91 | 1.48 | 3.0 | 1.25 | 0.82 | 3.87 | 3.79 |
#### Chart
| Category | | | | | |
|---|---|---|---|---|---|
| 24h | 0.0 | 0.0 | 0.0 | 0.0 | 0.0 |
| 48h | 1.14 | 2.91 | 2.64 | 1.48 | 3.67 |
| 72h | 1.99 | 4.03 | 4.16 | 2.88 | 4.08 |B)
Cluster 6
| Gene ID | 24h | 48h | 72h | Gen name |
| --- | --- | --- | --- | --- |
| BC2353 | 1.47 | 2.61 | 3.46 | Spo0A-P phosphatase |
| BC2142 | -0.32 | 2.59 | 3.71 | stage V sporulation protein S |
| BC1340 | -0.45 | 2.19 | 3.71 | Sporulation kinase KinE |
| BC2259 | 3.21 | 3.81 | 3.80 | Sporulation-control protein Spo0M |
| BC5482 | -3.31 | 0.36 | 0.77 | Sporulation initiation inhibitor protein soj |
Fig S4. A) Clustering of sporulation genes with STEM. Clusters 1-5 show sharp downregulation of most of sporulation genes. Cluster 6 contain five genes with an expression pattern of higher expression at 72 h than in previous strages. B) List of genes included in cluster 6 showing that three out of five genes are involved in negative regulation of sporulation and one gene involved in the last stage of spore maturation, proposed to play a role in the coat assembly. KinE, has been demonstrated low phosphorylation activity to the sporulation phosphorelay (Fujita and Losick, 2005).

### Slide 5
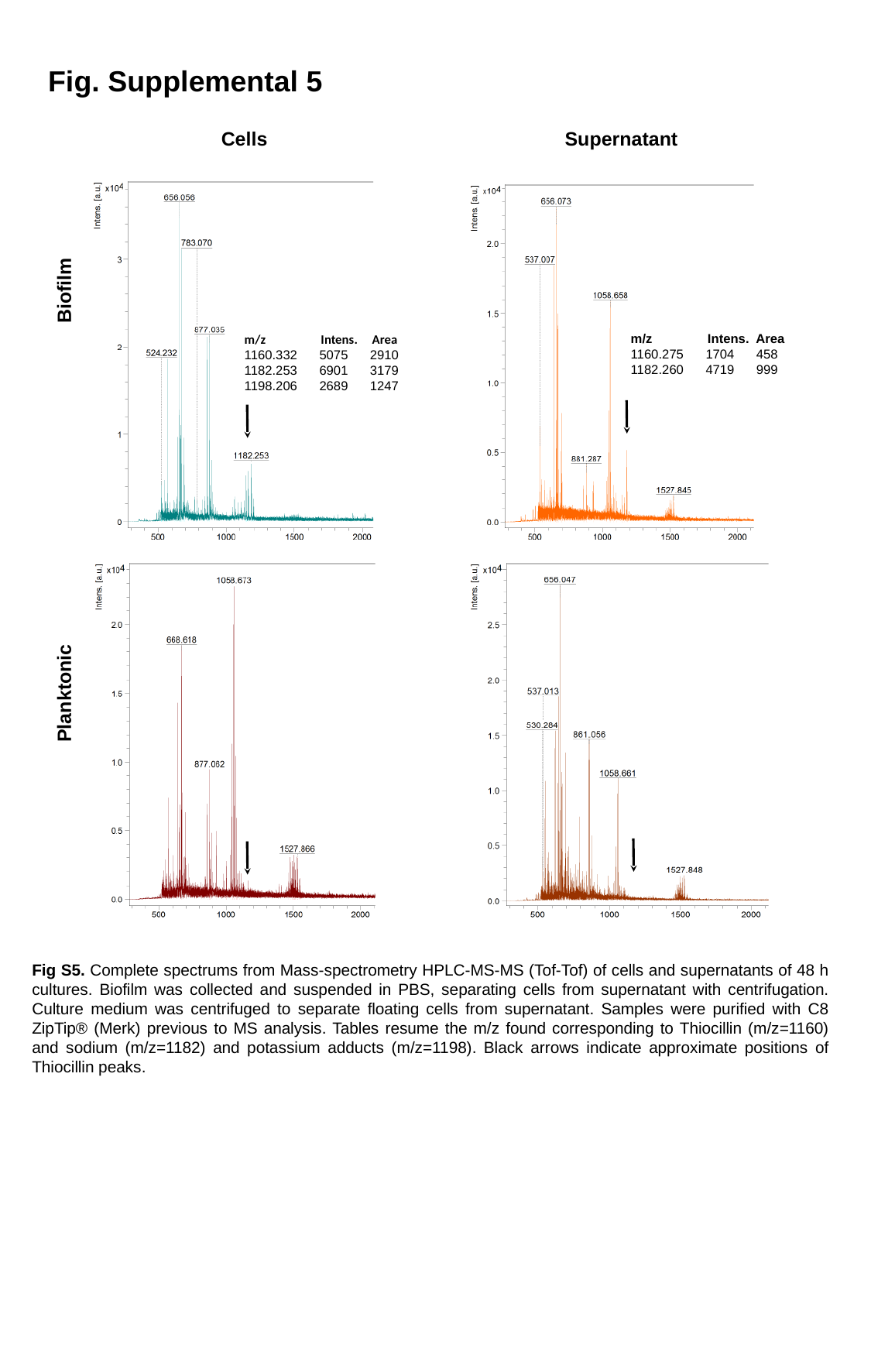

Fig. Supplemental 5
Supernatant
Cells
Biofilm
m/z Intens. Area
1160.275 1704 458
1182.260 4719 999
m/z Intens. Area
1160.332 5075 2910
1182.253 6901 3179
1198.206 2689 1247
Planktonic
Fig S5. Complete spectrums from Mass-spectrometry HPLC-MS-MS (Tof-Tof) of cells and supernatants of 48 h cultures. Biofilm was collected and suspended in PBS, separating cells from supernatant with centrifugation. Culture medium was centrifuged to separate floating cells from supernatant. Samples were purified with C8 ZipTip® (Merk) previous to MS analysis. Tables resume the m/z found corresponding to Thiocillin (m/z=1160) and sodium (m/z=1182) and potassium adducts (m/z=1198). Black arrows indicate approximate positions of Thiocillin peaks.

### Slide 6
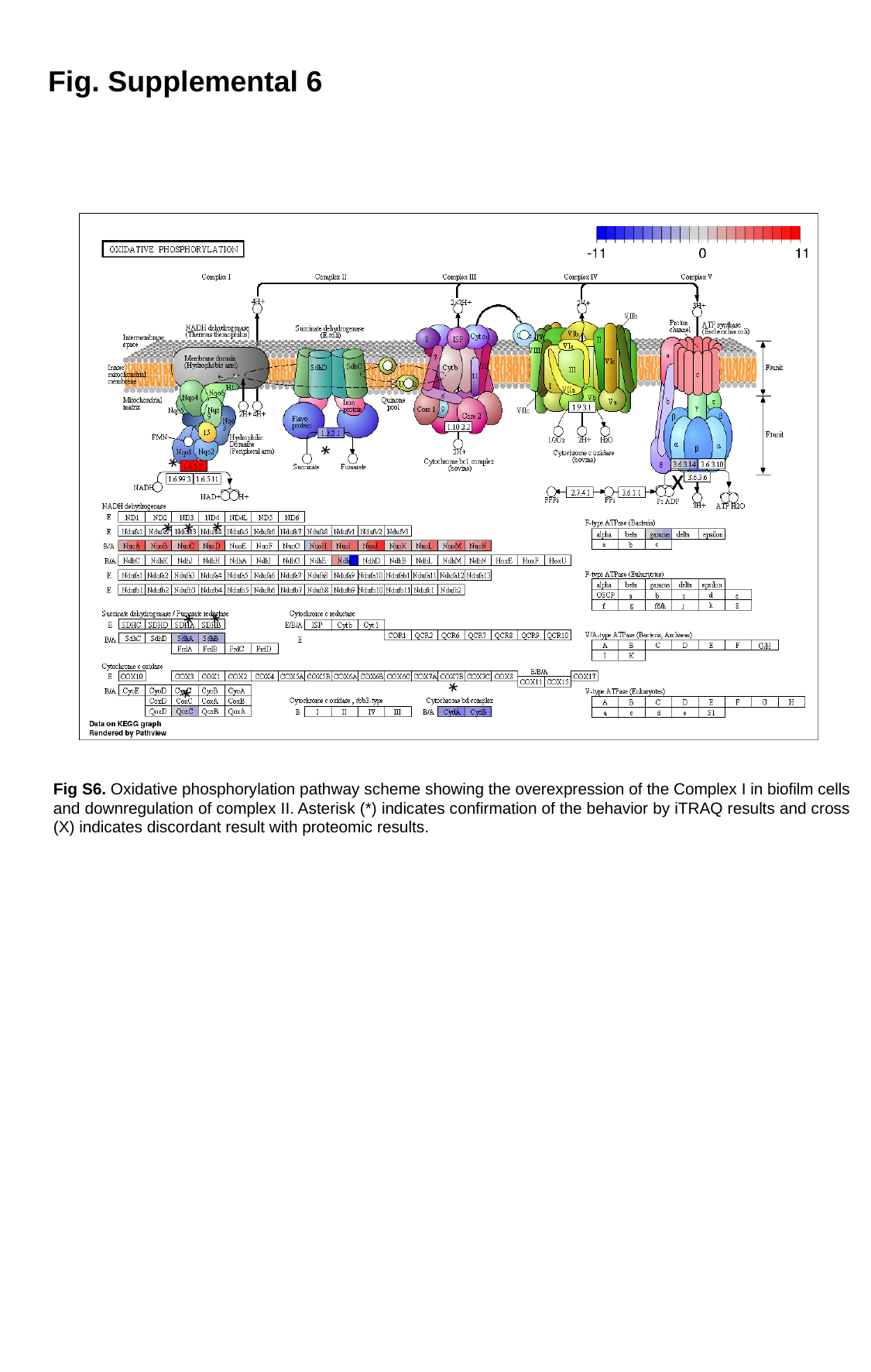

Fig. Supplemental 6
*
*
x
*
*
*
*
*
*
*
Fig S6. Oxidative phosphorylation pathway scheme showing the overexpression of the Complex I in biofilm cells and downregulation of complex II. Asterisk (*) indicates confirmation of the behavior by iTRAQ results and cross (X) indicates discordant result with proteomic results.

### Slide 7
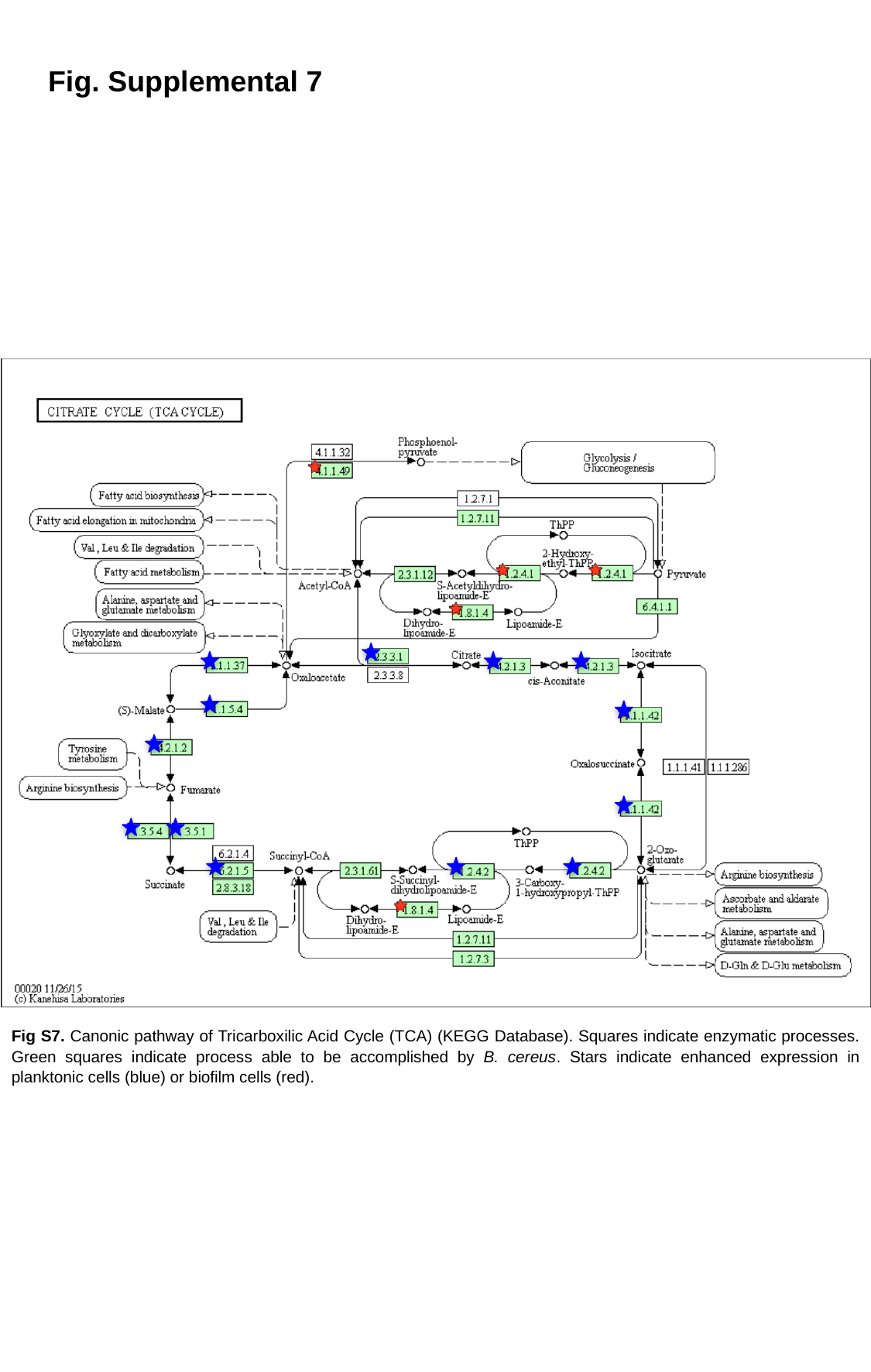

Fig. Supplemental 7
Fig S7. Canonic pathway of Tricarboxilic Acid Cycle (TCA) (KEGG Database). Squares indicate enzymatic processes. Green squares indicate process able to be accomplished by B. cereus. Stars indicate enhanced expression in planktonic cells (blue) or biofilm cells (red).

### Slide 8
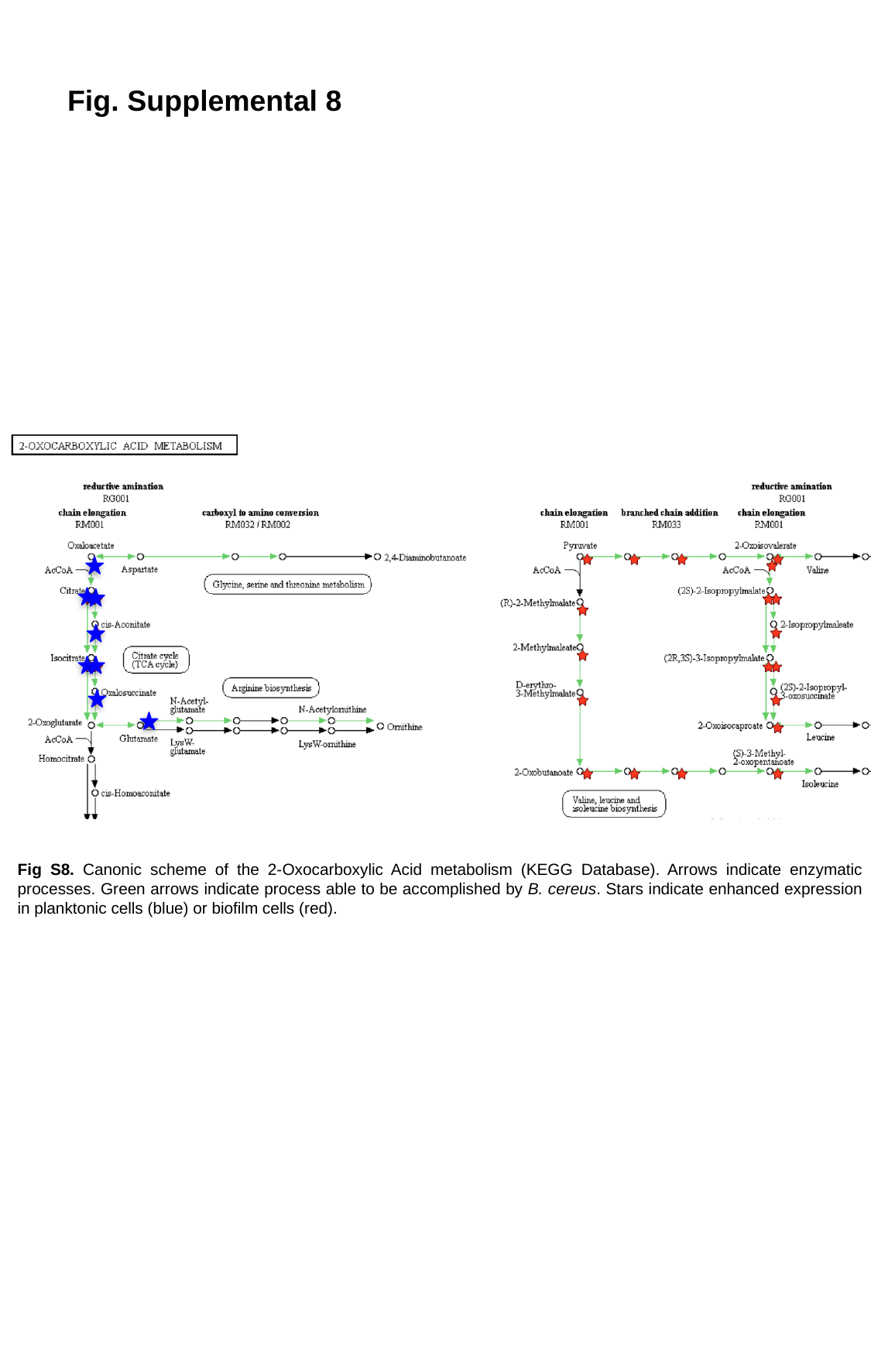

Fig. Supplemental 8
Fig S8. Canonic scheme of the 2-Oxocarboxylic Acid metabolism (KEGG Database). Arrows indicate enzymatic processes. Green arrows indicate process able to be accomplished by B. cereus. Stars indicate enhanced expression in planktonic cells (blue) or biofilm cells (red).

### Slide 9
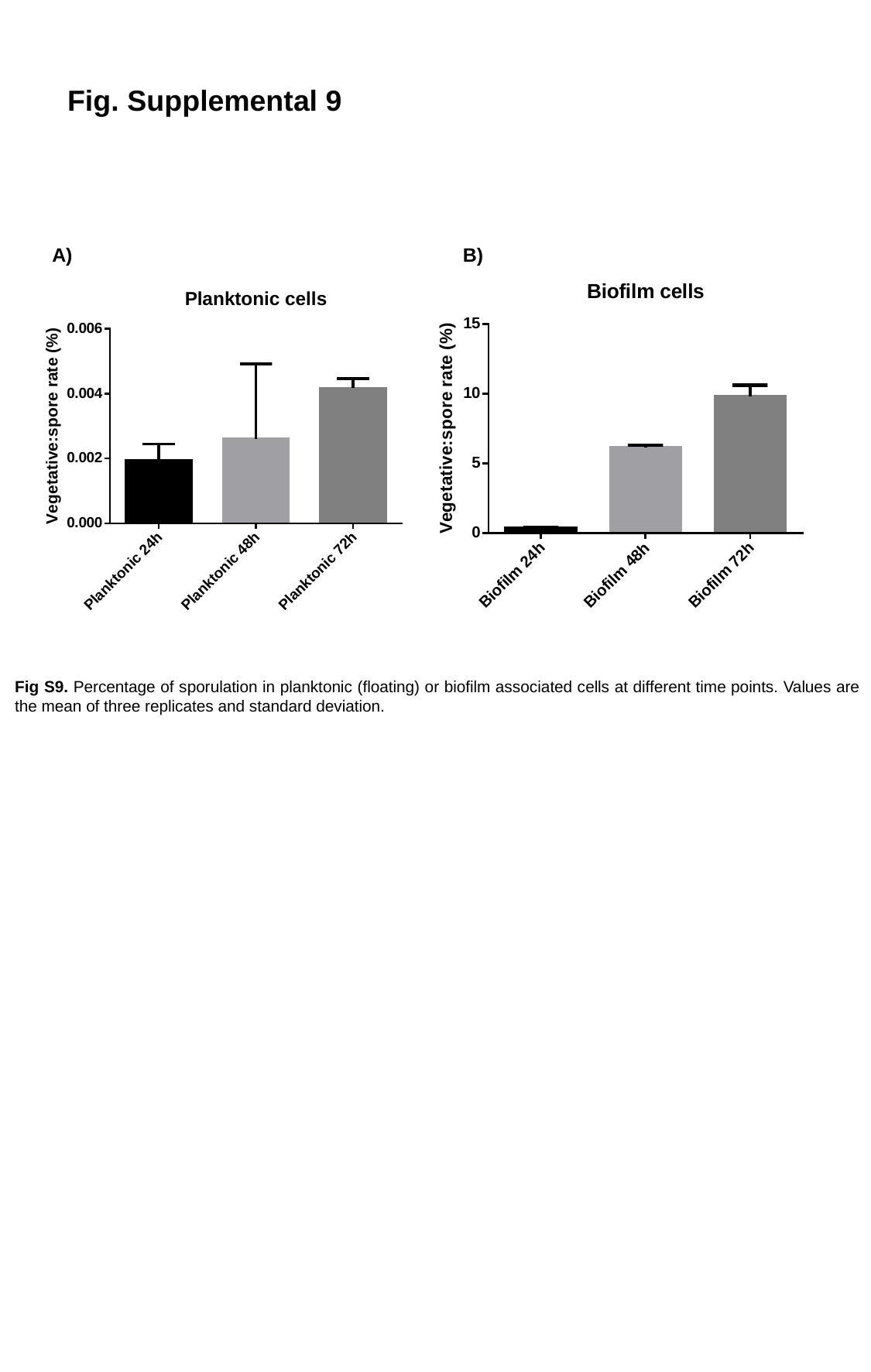

Fig. Supplemental 9
A)
B)
Fig S9. Percentage of sporulation in planktonic (floating) or biofilm associated cells at different time points. Values are the mean of three replicates and standard deviation.
