## Supplemental tables for "Biofilm formation displays intrinsic offensive and defensive features of *Bacillus cereus*"

**Table S1.** Expression pattern of genes involved in amino acid transport. Asterisk indicates confirmed behaviour at protein levels.

| | | Log2 (fold change) | | | |
|---|---|---|---|---|---|
| **Gene ID** | **24 h** | **48 h** | **72 h** | **Gene name** | **iTRAQ** |
| BC0401 | 1.38 | 4.05 | 4.71 | cystine transport system permease | * |
| BC0402 | 2.46 | 4.30 | 5.42 | cystine-binding protein | * |
| BC0403 | 1.24 | 4.12 | 4.70 | glutamine transport ATP-binding protein glnQ | * |
| BC0638 | 1.44 | 4.50 | 5.10 | Sodium/proton-dependent alanine carrier protein | * |
| BC0639 | 8.70 | 8.82 | 1.78 | glutamine transport ATP-binding protein glnQ | |
| BC0640 | 15.73 | 17.36 | 7.72 | glutamine-binding protein | * |
| BC0703 | 1.23 | 3.41 | 3.47 | Sodium/proline symporter | |
| BC0865 | 0.02 | -0.51 | -3.69 | arginine/ornithine antiporter | |
| BC1231 | -3.21 | -1.45 | -0.07 | Sodium/proline symporter | |
| BC1432 | -1.59 | -1.90 | -3.18 | proton/sodium-glutamate symport protein | * |
| BC1609 | 1.55 | 4.17 | 5.94 | Sodium/proline symporter | * |
| BC1927 | 2.11 | 5.79 | 6.66 | leu-, iso-, val-, trn-, ala-binding protein | * |
| BC2790 | 5.31 | 7.63 | 5.29 | glycine betaine transport system permease | |
| BC2980 | -0.14 | -0.53 | -4.31 | arginine permease | |
| BC3398 | 0.01 | -2.47 | -5.75 | Serine transporter | * |
| BC4071 | 2.82 | 5.62 | 6.82 | Sodium/proline symporter | * |
| BC4149 | -2.26 | -3.48 | -1.42 | arginine ABC transporter permease | * |
| BC4150 | -2.13 | -2.90 | -0.30 | arginine-binding protein | * |
| BC4242 | 1.12 | 2.20 | 4.00 | proton/sodium-glutamate symport protein | |
| BC5043 | -1.63 | -3.20 | -4.10 | Sodium/proton-dependent alanine carrier protein | |
| BC5051 | -3.53 | -1.60 | -3.76 | Sodium/proton-dependent alanine carrier protein | |
| BC5218 | -0.03 | -3.89 | -16.49 | proton/sodium-glutamate symport protein | |

**Table S2.** Expression pattern of genes involved in antimicrobial biosynthesis. Asterisk indicates confirmed behaviour at protein levels.

| | Log2 (fold change) | | | | |
|---|---|---|---|---|---|
| **Gene ID** | **24 h** | **48 h** | **72 h** | **Molecule** | **iTRAQ** |
| BC1201 | 2.87 | 5.63 | 1.75 | Tylosin | |
| BC1204 | 8.07 | 9.53 | 5.09 | Tylosin | |
| BC1206 | 7.01 | 8.13 | 2.73 | Tylosin | * |
| BC1208 | 5.00 | 6.74 | 1.04 | Streptomycin like | |
| BC1209 | 5.95 | 7.91 | 2.46 | Streptomycin like | * |
| BC1210 | 3.47 | 4.24 | -0.52 | Streptomycin like | |
| BC1211 | 3.78 | 6.19 | 2.59 | Streptomycin like | |
| BC1212 | 4.36 | 6.94 | 0.28 | Streptomycin like | * |
| BC1213 | 3.38 | 5.85 | -1.63 | Streptomycin like | |
| BC1214 | 3.10 | 5.17 | 0.92 | Streptomycin like | * |
| BC1248 | 3.77 | 5.88 | 2.80 | Bacteriocin | |
| BC1249 | 4.28 | 5.43 | 3.24 | Bacteriocin | |
| BC1250 | 4.31 | 5.00 | 1.78 | Bacteriocin | |
| BC2452 | 0.24 | 3.00 | 4.99 | Bacitracin | |
| BC2966 | 16.71 | 17.72 | 10.24 | Polyketide synthase | |
| BC5079 | -1.90 | 3.60 | 4.62 | Thiocillin | * |
| BC5080 | -1.58 | 3.00 | 3.75 | Thiocillin | * |
| BC5081 | -0.86 | 3.27 | 4.33 | Thiocillin | * |
| BC5082 | -2.43 | 3.20 | 4.32 | Thiocillin | * |
| BC5083 | -2.33 | 2.06 | 3.34 | Thiocillin | * |
| BC5084 | -1.76 | 2.48 | 3.69 | Thiocillin | * |
| BC5085 | -0.77 | 3.52 | 3.36 | Thiocillin | * |
| BC5086 | -1.76 | 2.83 | 4.04 | Thiocillin | * |
| BC5087 | 2.85 | 3.61 | 3.02 | Thiocillin | |
| BC5088 | 2.81 | 3.74 | 3.14 | Thiocillin | |
| BC5089 | 2.61 | 3.65 | 2.90 | Thiocillin | |
| BC5090 | 2.61 | 2.87 | 3.13 | Thiocillin | |
| BC3021 | 0.48 | 2.21 | 2.78 | Colicin | * |
| BC1426 | 2.01 | 8.10 | 8.50 | Porphirin | |
| BC1427 | 1.46 | 5.19 | 4.77 | Porphirin | |
| BC1428 | 3.04 | 6.66 | 6.96 | Porphirin | * |
| BC2133 | -2.57 | -4.22 | -11.87 | Porphirin | |
| BC2134 | -2.99 | -4.05 | -6.25 | Porphirin | |
| BC4468 | 2.84 | 3.27 | 3.62 | Porphirin | |

**Table S3.** Specialization of biofilm and floating cells in distinct but coordinated metabolic activities. Genes belonging to cluster 1.

| GeneIDs | Planktonic(blue)/Biofilm(red) | DAVID Functional annotation |
|---|---|---|
| BC0344 | blue | 1-pyrroline-5-carboxylate dehydrogenase |
| BC0410 | blue | Crp family transcriptional regulator |
| BC0411 | blue | hypothetical protein |
| BC0466 | blue | fumarate hydratase |
| BC0491 | blue | formate acetyltransferase |
| BC0492 | blue | pyruvate formate-lyase activating enzyme |
| BC0589 | blue | formate dehydrogenase alpha chain |
| BC0590 | blue | hypothetical protein |
| BC0611 | blue | aspartate ammonia-lyase |
| BC0612 | blue | L-lactate permease |
| BC0621 | blue | 2-amino-3-ketobutyrate CoA ligase |
| BC0849 | blue | acetyltransferase |
| BC0898 | blue | enoyl-CoA hydratase |
| BC1149 | blue | ornithine--oxo-acid transaminase |
| BC1231 | blue | sodium/proline symporter |
| BC1252 | blue | 2-oxoglutarate dehydrogenase subunit E1 |
| BC1746 | blue | asparagine synthetase AsnA |
| BC2220 | blue | alcohol dehydrogenase |
| BC2758 | blue | metal-dependent hydrolase |
| BC2896 | blue | aspartate aminotransferase |
| BC2959 | blue | malate:quinone oxidoreductase |
| BC3616 | blue | aconitate hydratase |
| BC3650 | blue | imidazolonepropionase |
| BC3651 | blue | urocanate hydratase |
| BC3652 | blue | histidine ammonia-lyase |
| BC3653 | blue | anti-terminator HutP |
| BC3833 | blue | succinyl-CoA synthetase subunit alpha |
| BC3834 | blue | succinyl-CoA synthetase subunit beta |
| BC4023 | blue | acetyl-CoA acetyltransferase |
| BC4224 | blue | glycine dehydrogenase subunit 2 |
| BC4225 | blue | glycine dehydrogenase subunit 1 |
| BC4226 | blue | glycine cleavage system aminomethyltransferase T |
| BC4365 | blue | bifunctional acetaldehyde-CoA/alcohol dehydrogenase |
| BC4516 | blue | succinate dehydrogenase iron-sulfur subunit |
| BC4517 | blue | succinate dehydrogenase flavoprotein subunit |
| BC4592 | blue | malate dehydrogenase |
| BC4593 | blue | isocitrate dehydrogenase |
| BC4594 | blue | citrate synthase |
| BC4870 | blue | L-lactate dehydrogenase |

| | | |
|---|---|---|
| BC4995 | blue | regulatory protein |
| BC4996 | blue | L-lactate dehydrogenase |
| BC5002 | blue | acyl-CoA dehydrogenase |
| BC5003 | blue | acetyl-CoA acetyltransferase |
| BC5004 | blue | enoyl-CoA hydratase |
| BC5006 | blue | Prolyne dehydrogenase |
| BC5228 | blue | L-lactate permease |
| BC5342 | blue | acyl-CoA dehydrogenase |
| BC5344 | blue | acetyl-CoA acetyltransferase |
| BC5345 | blue | Iron-sulphur-binding reductase |
| BC5438 | blue | antiholin-like protein LrgB |
| BC5439 | blue | murein hydrolase regulator LrgA |
| BC0355 | red | 4-aminobutyrate--2-oxoglutarate transaminase |
| BC0356 | red | sigma-54-dependent transcriptional activator |
| BC0357 | red | succinate-semialdehyde dehydrogenase [NADP+] |
| BC1036 | red | glycerol-3-phosphate dehydrogenase |
| BC1301 | red | two component system histidine kinase |
| BC1396 | red | branched-chain amino acid aminotransferase |
| BC1398 | red | acetolactate synthase small subunit |
| BC1399 | red | ketol-acid reductoisomerase |
| BC1400 | red | 2-isopropylmalate synthase |
| BC1401 | red | 3-isopropylmalate dehydrogenase |
| BC1402 | red | 3-isopropylmalate dehydratase large subunit |
| BC1610 | red | aminotransferase |
| BC1611 | red | hypothetical protein |
| BC1776 | red | branched-chain amino acid aminotransferase |
| BC1777 | red | acetolactate synthase 3 catalytic subunit |
| BC2285 | red | citrate synthase 3 |
| BC2286 | red | 2-methylcitrate dehydratase |
| BC2287 | red | methylisocitrate lyase |
| BC2288 | red | acyl-CoA dehydrogenase |
| BC2289 | red | 3-hydroxyisobutyrate dehydrogenase |
| BC2290 | red | methylmalonate-semialdehyde dehydrogenase |
| BC2292 | red | 3-hydroxyisobutyryl-CoA hydrolase |
| BC2776 | red | dihydrolipoamide dehydrogenase |
| BC2778 | red | acetoin dehydrogenase E1 component beta-subunit |
| BC3555 | red | aldehyde dehydrogenase |
| BC3939 | red | formamidase |
| BC4325 | red | late competence protein ComER |
| BC4595 | red | hypothetical protein |
| BC4644 | red | PhnB protein |
| BC4645 | red | acetyl-coenzyme A synthetase |
| BC4762 | red | phosphoenolpyruvate carboxykinase |
| BC5133 | red | holin-like protein |

**Table S4.** KEGG pathways enrichment data for ALL, blue (planktonic), and red (biofilm) genes in cluster 1. Significance tests p-value and Benjamini are shown, and the number and percentage of gene over the total genes in Cluster 1 is also shown for each KEGG pathway.

| KEGG: Metabolic pathways | P-Value | Benjamini | #genes | %genes |
|---|---|---|---|---|
| ALL | 8.00E-10 | 6.80E-09 | 52 | 62.7 |
| BLUE (Planktonic) | 3.50E-07 | 2.10E-06 | 34 | 66.7 |
| RED (Biofilm) | 1.70E-03 | 6.10E-03 | 18 | 56.2 |

| KEGG: Biosynthesis of antibiotics | P-Value | Benjamini | #genes | %genes |
|---|---|---|---|---|
| ALL | 5.80E-14 | 2.90E-12 | 35 | 42.2 |
| BLUE (Planktonic) | 1.60E-10 | 6.60E-09 | 24 | 47.1 |
| RED (Biofilm) | 5.00E-04 | 2.30E-03 | 11 | 34.4 |

| KEGG: Biosynthesis of secondary metabolites | P-Value | Benjamini | #genes | %genes |
|---|---|---|---|---|
| ALL | 8.90E-13 | 2.30E-11 | 40 | 48.2 |
| BLUE (Planktonic) | 7.10E-08 | 5.90E-07 | 25 | 49 |
| RED (Biofilm) | 1.90E-05 | 1.70E-04 | 15 | 46.9 |

| KEGG: Citrate cycle (TCA cycle) | P-Value | Benjamini | #genes | %genes |
|---|---|---|---|---|
| ALL | 9.00E-13 | 1.50E-11 | 15 | 18.1 |
| BLUE (Planktonic) | 1.00E-09 | 1.40E-08 | 11 | 21.6 |
| RED (Biofilm) | 8.70E-03 | 2.10E-02 | 4 | 12.5 |

| KEGG: 2-Oxocarboxylic acid metabolism | P-Value | Benjamini | #genes | %genes |
|---|---|---|---|---|
| ALL | 2.80E-10 | 2.80E-09 | 13 | 15.7 |
| BLUE (Planktonic) | 4.00E-02 | 1.00E-01 | 4 | 7.8 |
| RED (Biofilm) | 1.70E-09 | 3.20E-08 | 9 | 28.1 |

| KEGG: Glyoxylate and dicarboxylate metabolism | P-Value | Benjamini | #genes | %genes |
|---|---|---|---|---|

| | | | | |
|---|---|---|---|---|
| ALL | 4.10E-09 | 2.60E-08 | 13 | 15.7 |
| BLUE (Planktonic) | 1.10E-07 | 7.60E-07 | 10 | 19.6 |
| RED (Biofilm) | 9.30E-02 | 1.90E-01 | 3 | 9.4 |

| **KEGG: Pyruvate metabolism** | **P-Value** | **Benjamini** | **#genes** | **%genes** |
|---|---|---|---|---|
| ALL | 2.20E-08 | 1.20E-07 | 15 | 18.1 |
| BLUE (Planktonic) | 7.50E-05 | 3.40E-04 | 9 | 17.6 |
| RED (Biofilm) | 1.10E-03 | 4.40E-03 | 6 | 18.8 |

| **KEGG: Propanoate metabolism** | **P-Value** | **Benjamini** | **#genes** | **%genes** |
|---|---|---|---|---|
| ALL | 5.50E-08 | 2.80E-07 | 11 | 13.3 |
| BLUE (Planktonic) | 5.70E-05 | 2.90E-04 | 7 | 13.7 |
| RED (Biofilm) | 7.10E-03 | 2.00E-02 | 4 | 12.5 |

| **KEGG: Butanoate metabolism** | **P-Value** | **Benjamini** | **#genes** | **%genes** |
|---|---|---|---|---|
| ALL | 1.20E-07 | 5.20E-07 | 11 | 13.3 |
| BLUE (Planktonic) | 8.90E-05 | 3.60E-04 | 7 | 13.7 |
| RED (Biofilm) | 8.70E-03 | 2.10E-02 | 4 | 12.5 |

**Table S5**. Gene composition of three general enriched KEGG pathways in Cluster 1 is indicated by "x" symbol.

| KEGG: Metabolic pathways | KEGG: synthesis of secondary metabolites | KEGG: synthesis of antibiotics | Functional annotation | Gene |
|---|---|---|---|---|
| x | | | 1-Pyrroline-5-Carboxylate dehydrogenase | BC0344 |
| x | | | 4-Aminobutyrate--2-Oxoglutarate transaminase | BC0355 |
| | x | | Glycerol-3-Phosphate dehydrogenase | BC1036 |
| x | | | Succinate-Semialdehyde dehydrogenase [NADP+] | BC0357 |
| x | x | x | Fumarate hydratase | BC0466 |
| x | | | Formate acetyltransferase | BC0491 |
| x | | | formate dehydrogenasealpha chain | BC0589 |
| x | | | aspartate ammonia-lyase | aspA |
| x | x | x | acetyltransferase | BC0849 |
| x | x | x | ornithine--oxo-acid transaminase | rocD |
| x | x | x | 2-oxoglutarate dehydrogenasesubunit E1 | sucA |
| x | x | x | branched-chain aminoacid aminotransferase | BC1396 |
| x | x | x | acetolactate synthasesmall subunit | ilvH |
| x | x | x | ketol-acid reductoisomerase | BC1399 |
| x | x | | 2-isopropylmalate synthase | BC1400 |
| x | x | | 3-isopropylmalate dehydrogenase | BC1401 |
| x | x | | 3-isopropylmalate dehydrataselarge subunit | BC1402 |
| x | x | | asparagine synthetaseAsnA | BC1746 |
| x | x | x | branched-chain aminoacid aminotransferase | BC1776 |
| x | x | x | acetolactate synthase small subunit | BC1777 |
| x | x | x | alcohol dehydrogenase | BC2220 |
| x | x | x | citrate synthase3 | BC2285 |
| x | | | 3-hydroxyisobutyrate dehydrogenase | BC2289 |
| x | | | methylmalonate-semialdehyde dehydrogenase | acylating |
| x | x | | metal-dependent hydrolase | BC2758 |
| x | x | x | dihydrolipoamide dehydrogenase | acoL |
| x | x | x | acetoin dehydrogenase E1 component beta-subunit | BC2778 |
| x | x | x | malate:quinone oxidoreductase | BC2959 |
| x | x | x | aldehyde dehydrogenase | BC3555 |
| x | x | x | aconitate hydratase | BC3616 |
| x | x | | imidazolonepropionase | BC3650 |
| x | x | | urocanate hydratase | BC3651 |
| x | x | | histidine ammonia-lyase | hutH |
| x | x | x | succinyl-CoA synthetasesubunit alpha | BC3833 |
| x | | x | succinyl-CoA synthetasesubunit beta | sucC |
| x | | x | acetyl-CoA acetyltransferase | BC4023 |
| x | | x | glycine dehydrogenasesubunit 2 | BC4224 |

| | | | | |
|---|---|---|---|---|
| x | x | x | glycine dehydrogenasesubunit 1 | BC4225 |
| x | x | x | glycine cleavage system aminomethyltransferase T | gcvT |
| x | x | x | bifunctional acetaldehyde-CoA/alcoholdehydrogenase | BC4365 |
| x | x | x | succinate dehydrogenaseiron-sulfur subunit | sdhB |
| x | x | x | succinate dehydrogenaseflavoprotein subunit | sdhA |
| x | x | x | malate dehydrogenase | BC4592 |
| x | x | x | isocitrate dehydrogenase | BC4593 |
| x | x | x | citrate synthase | BC4594 |
| x | x | x | acetyl-coenzyme Asynthetase | BC4645 |
| x | x | x | phosphoenolpyruvate carboxykinase | BC4762 |
| x | x | x | L-lactate dehydrogenase | ldh |
| x | x | x | L-lactate dehydrogenase | ldh |
| x | | x | acetyl-CoA acetyltransferase | BC5003 |
| x | x | | enoyl-CoA hydratase | BC5004 |
| x | x | x | Prolyne dehydrogenase | BC5006 |
| x | | x | acetyl-CoA acetyltransferase | BC5344 |

**Table S6**. Specialization of biofilm and floating cells in distinct but coordinated metabolic activities. Genes belonging to cluster 2.

| GeneIDs | Planktonic(blue)/Biofilm(red) | DAVID functional annotation |
|---|---|---|
| BC0049 | red | SspF protein |
| BC0069 | red | stage II sporulation protein E |
| BC0169 | red | spore germination protein GerD |
| BC0170 | blue | #N/A |
| BC0263 | red | #N/A |
| BC0520 | blue | #N/A |
| BC0603 | red | hypothetical protein |
| BC0875 | red | small acid-soluble spore protein |
| BC0877 | red | hypothetical protein |
| BC1012 | blue | #N/A |
| BC1225 | blue | #N/A |
| BC1500 | red | hypothetical protein |
| BC1501 | red | hypothetical protein |
| BC1502 | red | hypothetical protein |
| BC1509 | red | stage IV sporulation protein A |
| BC1520 | red | #N/A |
| BC1984 | red | small acid-soluble spore protein |
| BC1994 | blue | #N/A |
| BC2010 | red | #N/A |
| BC2050 | red | #N/A |
| BC2095 | red | hypothetical protein |
| BC2536 | red | cell wall hydrolase cwlJ |
| BC3106 | red | small acid-soluble spore protein |
| BC3605 | red | #N/A |
| BC3770 | red | spore coat protein E |
| BC3783 | red | #N/A |
| BC3800 | red | dipicolinate synthase subunit B |
| BC3801 | red | dipicolinate synthase subunit A |
| BC3802 | red | hypothetical protein |
| BC3902 | red | hypothetical protein |
| BC3905 | red | sporulation sigma-E factor processing peptidase |
| BC3922 | red | prespore specific transcriptional activator rsfA |
| BC4067 | red | stage V sporulation protein AD |
| BC4073 | red | anti-sigma F factor |
| BC4088 | red | hypothetical protein |
| BC4186 | red | stage III sporulation protein AH |
| BC4187 | red | stage III sporulation protein AG |
| BC4188 | red | stage III sporulation protein AF |
| BC4191 | red | stage III sporulation protein AC |
| BC4192 | red | stage III sporulation protein SpoAB |
| BC4193 | red | #N/A |

| | | |
|---|---|---|
| BC4194 | red | hypothetical protein |
| BC4288 | blue | membrane-attached cytochrome c550 |
| BC4440 | red | #N/A |
| BC4466 | red | CotS-related protein |
| BC4467 | red | stage VI sporulation protein D |
| BC4495 | red | germination protein germ |
| BC4563 | red | small acid-soluble spore protein SspI |
| BC4577 | red | hypothetical protein |
| BC4606 | red | hypothetical protein |
| BC4640 | red | hypothetical protein |
| BC4641 | red | hypothetical protein |
| BC4642 | blue | inorganic polyphosphate/ATP-NAD kinase |
| BC4660 | blue | #N/A |
| BC4662 | blue | #N/A |
| BC4899 | red | hypothetical protein |
| BC4923 | blue | #N/A |
| BC4924 | blue | #N/A |
| BC5145 | red | hypothetical protein |
| BC5147 | red | stage V sporulation protein AC |
| BC5148 | red | stage V sporulation protein AD |
| BC5149 | red | stage V sporulation protein AE |
| BC5282 | red | stage III sporulation protein D |
| BC5283 | red | stage II sporulation protein Q |
| BC5287 | red | stage II sporulation protein D |
| BC5289 | red | #N/A |
| BC5385 | red | prespore specific transcriptional activator rsfA |
| BC5390 | red | cell wall hydrolase cwlJ |
| BC5480 | red | hypothetical protein |
| BC5391 | red | hypothetical protein |
| BC4607 | red | hypothetical protein |

**Table S7.** Specialization of biofilm and floating cells in distinct but coordinated metabolic activities. Genes belonging to cluster 3.

| GeneIDs | Planktonic(blue)/Biofilm(red) | DAVID functional annotation |
|---|---|---|
| BC0114 | blue | RNA polymerase factor sigma-70 |
| BC0404 | blue | methyl-accepting chemotaxis protein |
| BC0405 | blue | #N/A |
| BC0422 | blue | methyl-accepting chemotaxis protein |
| BC0559 | blue | methyl-accepting chemotaxis protein |
| BC0576 | blue | methyl-accepting chemotaxis protein |
| BC0678 | blue | methyl-accepting chemotaxis protein |
| BC0679 | red | #N/A |
| BC1625 | blue | flagellar motor protein MotP |
| BC1626 | blue | #N/A |
| BC1627 | blue | chemotaxis protein CheY |
| BC1628 | blue | chemotaxis protein CheA |
| BC1629 | blue | flagellar motor switch protein |
| BC1630 | blue | hypothetical protein |
| BC1636 | blue | flagellar hook-associated protein FlgK |
| BC1637 | blue | flagellar hook-associated protein FlgL |
| BC1638 | blue | flagellar capping protein |
| BC1639 | blue | flagellar protein fliS |
| BC1643 | blue | flagellar hook-basal body protein FliE |
| BC1644 | blue | flagellar MS-ring protein |
| BC1645 | blue | flagellar motor switch protein G |
| BC1646 | blue | flagellar assembly protein H |
| BC1647 | blue | flagellum-specific ATP synthase |
| BC1651 | blue | flagellar hook protein FlgE |
| BC1653 | blue | hypothetical protein |
| BC1654 | blue | chemotaxis protein CheV |
| BC1657 | blue | flagellin |
| BC1658 | blue | flagellin |
| BC1659 | blue | flagellin |
| BC1660 | blue | soluble lytic murein transglycosylase |
| BC1662 | blue | flagellar motor switch protein FliM |
| BC1664 | blue | flagellar motor switch protein fliN |
| BC1671 | blue | flagellar basal body rod protein FlgG |
| BC1672 | blue | metal-dependent hydrolase related to alanyl-tRNA synthetase |
| BC2006 | blue | methyl-accepting chemotaxis protein |
| BC2766 | blue | #N/A |
| BC3903 | red | sporulation sigma factor SigG |
| BC3904 | red | sporulation sigma factor SigE |
| BC4071 | red | Sodium/proline symporter |
| BC4072 | red | sporulation sigma factor SigF |
| BC4074 | red | anti-sigma F factor antagonist |
| BC4512 | blue | flagellar motor protein MotB |

| | | |
|---|---|---|
| BC4513 | blue | flagellar motor protein MotA |
| BC5009 | blue | methyl-accepting chemotaxis protein |
| BC5034 | blue | methyl-accepting chemotaxis protein |
| BC5065 | blue | #N/A |
| BC5143 | blue | #N/A |
| BC1631 | blue | hypothetical protein |
| BC1640 | blue | hypothetical protein |
| BC1648 | blue | cytoplasmic protein |
| BC1652 | blue | hypothetical protein |
| BC5035 | red | #N/A |
| BC5066 | blue | endonuclease/exonuclease/phosphatase family protein |
